# Transcriptional logic of cell fate specification and axon guidance in early born retinal neurons revealed by single-cell mRNA profiling

**DOI:** 10.1101/497081

**Authors:** Quentin Lo Giudice, Marion Leleu, Pierre J. Fabre

**Affiliations:** Department of Basic Neurosciences, University of Geneva, 1205 Geneva, Switzerland.; School of Life Sciences, Ecole Polytechnique Fédérale, Lausanne, 1015 Lausanne, Switzerland.

## Abstract

Retinal ganglion cells (RGC), together with cone photoreceptors, horizontal cells (HC) and amacrine cells (AC), are the first classes of neurons produced in the retina. Here we have profiled 5348 single retinal cells and provided a comprehensive transcriptomic atlas showing the broad diversity of the developing retina at the time when the four early-born cells are being produced. Our results show the transcriptional sequences that establish the hierarchical ordering of early cell fate specification in the retina. RGC maturation follows six waves of gene expression, giving new insight into the regulatory logic of RGC differentiation. Early-generated RGCs transcribe an increasing amount of guidance cues for young peripheral RGC axons that express the matching receptors. Finally, spatial signatures in sub-populations of RGCs allowed to define novel molecular markers that are spatially restricted during the development of the retina. Altogether this study is a valuable resource that identifies new players in mouse retinal development, shedding light on transcription factors sequence and guidance cues dynamics in space and time.

## INTRODUCTION

Understanding how diverse neuronal cell types emerge in the mammalian central nervous system (CNS) is essential to our understanding of the logic of neuronal network assembly. The timing of production of the different classes of neurons, and later glial cells, is thought to be crucial to the establishment of functionally efficient networks (Rossi et al., 2017). Interestingly, recent technical progress, such as single-cell RNA-seq, has allowed the identification of an increasing number of cell types (Poulin et al., 2016; Tasic et al., 2016; Yuan et al., 2018; Zeisel et al., 2015), but untangling the transcriptional features involved in the chronological generation of diverse classes of neuron and glial cells remains a conundrum. The mouse retina was one of the first CNS tissues analyzed using single-cell RNA-seq (Macosko et al., 2015; Shekhar et al., 2016; Trimarchi et al., 2008). The retina also represents an excellent model to decipher how diverse neuronal types are produced, as it is one of the simpler parts of the central nervous system with only six classes of neurons that have been extensively characterized both molecularly and morphologically (Cherry et al., 2009; Livesey and Cepko, 2001; Sanes and Masland, 2015). The retina produces its different classes of neurons in two waves (Rapaport et al., 2004). The first gives rise to the early-born cell types from embryonic days (E) 10 to 17, which comprise the retinal ganglion cells (RGC), the horizontal cells (HC), the amacrine cells (AC) and the cone photoreceptors (cones). In the second wave, AC cells are still produced, together with bipolar cells (BC) and rod photoreceptors, from E14 to post-natal day 5. Recently, bipolar cells were shown to exhibit an unexpected level of sub-classes in the adult mouse, which were associated with morphological features, further showing that the use of single-cell RNA-seq can unveil novel transcriptional programs that match the differentiation of these late-born retinal cells (Shekhar et al., 2016). Also very diverse, with more than 30 subtypes based on their dendritic morphologies, the RGCs are early-born cells that represent the sole output from the retina to the brain, where they can follow many different trajectories, reaching up to 46 targets (Martersteck et al., 2017; Morin and Studholme, 2014; Rivlin-Etzion et al., 2011; Seabrook et al., 2017). Recent work in adult and juvenile RGCs revealed a high degree of transcriptional heterogeneity (Macosko et al., 2015; Rheaume et al., 2018). However, the manner in which retinal progenitors give rise to this extreme heterogeneity is not fully understood.

Seminal work using vector lineage tracing identified some fundamental aspects of the logic that allows various retinal cell fates (Turner and Cepko, 1987; Turner et al., 1990; Wetts and Fraser, 1988; Wetts et al., 1989). This process involves intrinsic components, with sequential expression of transcription factors playing a key role in driving competence in progenitors to generate distinct cell types (Cayouette et al., 2003; Cepko, 2014). Recent advances in single-cell genomics have proven the capacity to delineate neuronal lineages using either transcriptomic profiles (single-cell RNA-seq) or enhancer signatures (ATAC-seq) (Kester and van Oudenaarden, 2018).

Here we used single-cell transcriptomic reconstructions to unveil the programs at work in the early specification of mouse retina. Taking advantage of the temporal organization of the retina, we identified couples of ligand-receptors as putative axon guidance pairs that can guide RGC-growing axons on their paths to exit the retina. Finally, exploiting the spatial distribution of RGC cells of the retina, we were able to attribute dorso-ventral and temporo-nasal scores to each RGC. This last refinement allowed us to identify RGCs from the ventro-temporal crescent of the retina, in which we found a significant subset of cells expressing ipsilateral-projecting RGC genes. This last analysis revealed genes that appear enriched to this subset of RGCs, giving new insight into the temporal development of axon guidance transcriptional programs.

## RESULTS

The goal of this study is threefold: first, to trace the origin of early-born retinal fates; second, to infer the spatial relationships across retinal neurons, and third to identify in the RGC group the transcriptional signature linked with projections through different routes in the developing brain.

### Cell type identification

To address our first point, we produced the transcriptional profiles of early-born retinal progenitors obtained from 5348 single-cells of E15.5 retina using 10X genomics (**Fig. 1A-B**). Cells were distributed in 14 clusters in a t-SNE obtained from merging two replicates (**Fig. 1C**). Both replicates were preliminary analyzed separately using the chromium v2 analysis software (Cell Ranger) giving similar clustering based on the t-SNE of 2675 and 2673 cells each (**Fig. S1A-B**). Each cluster was characterized using known marker genes (**Figs. 1D-I and S2**) (Bassett and Wallace, 2012). The central clusters (0–3) were composed of retinal cycling cells that we refer to as the retinal progenitor cells (RPC) and expressing *Sox2, Fos* and *Hes1* (**Fig. 1D**). Emanating from this core unit we could see a narrower group (cluster 4) containing cells expressing the cell cycle exit genes *Top2a, Prc1 as* well as neuronal-specific genes *Sstr2, Penk* and *Btg2* (**Figs. 1E and S2A-B**). The latter gene was previously shown to induce neuronal differentiation (el-Ghissassi et al., 2002) and was also identified in retinal progenitor cells (Trimarchi et al., 2008). The progression from these cycling cells led to a cluster aggregating early neuroblast transcription factors including *Neurod4* and *Pax6* (Neuroblast, cluster 5) as well as genes known for their importance in the initiation of axonal growth (*Pcdh17*) (**Fig. 1E**, **Fig. S2C-E**) (Hayashi et al., 2014). Following the neuroblast bottleneck furrow we observed three distinct branches. The main one (clusters 9–12) was characterized by RGC markers (*Isl1, Pou4f2, Pou6f2* and *Elavl4*) (**Figs. 1F**, **S2F-G**). The second neuronal group (clusters 7–8) was disconnected from the others and composed of both AC in the root part (*Onecut2*+*, Prox1*+) and ends up with HC (*Onecut*1+, *Lhx1*+) (**Fig. 1G**, **Fig. S2H-I**). The last neuronal branch (cluster 6) was positive for *Otx2, Crx* and the early cone marker *Thrb*, a signature for cone photoreceptors (**Fig. 1H, Fig. S2J-K**). The most excluded cluster was positive for several RGC genes, as well as enriched in mitochondrial genes (cluster 13) (**Table S1**). Considering its proportion and its content (e.g. *Hb* genes), we attributed the possible identity of the ciliary bodies (CB) to this group of cells. The main neuronal clusters were then validated with *in situ* hybridization (ISH) (**Fig. S3A-G**). In the cone cluster, we found that, in addition to *Thrb* and *Crx, Rbp4* transcripts were highly enriched. We validated the specific expression of *Rbp4* using mice expressing *Cre* under the control of the *Rbp4* promoter (**Fig. 1I**). Both the morphology and the positions of *Rbp4* positive cells followed cone hallmarks, as at this stage they already have a distinctive morphology and are aligned along the apical side of the retina (Decembrini et al., 2017). Finally, we represented these 14 clusters on a heatmap where we show the top 10 differentially expressed genes (**Fig. 1J**, top 15 in **Table S1**). This balanced clustering respected neuronal types similitudes and highlighted the major transcriptional modules that define the differentiation process into early retinal cell types (**Fig. 1K**). Our analysis provides a comprehensive characterization of the main groups of cells, allowing further exploration of each cell type.

**Figure 1.**
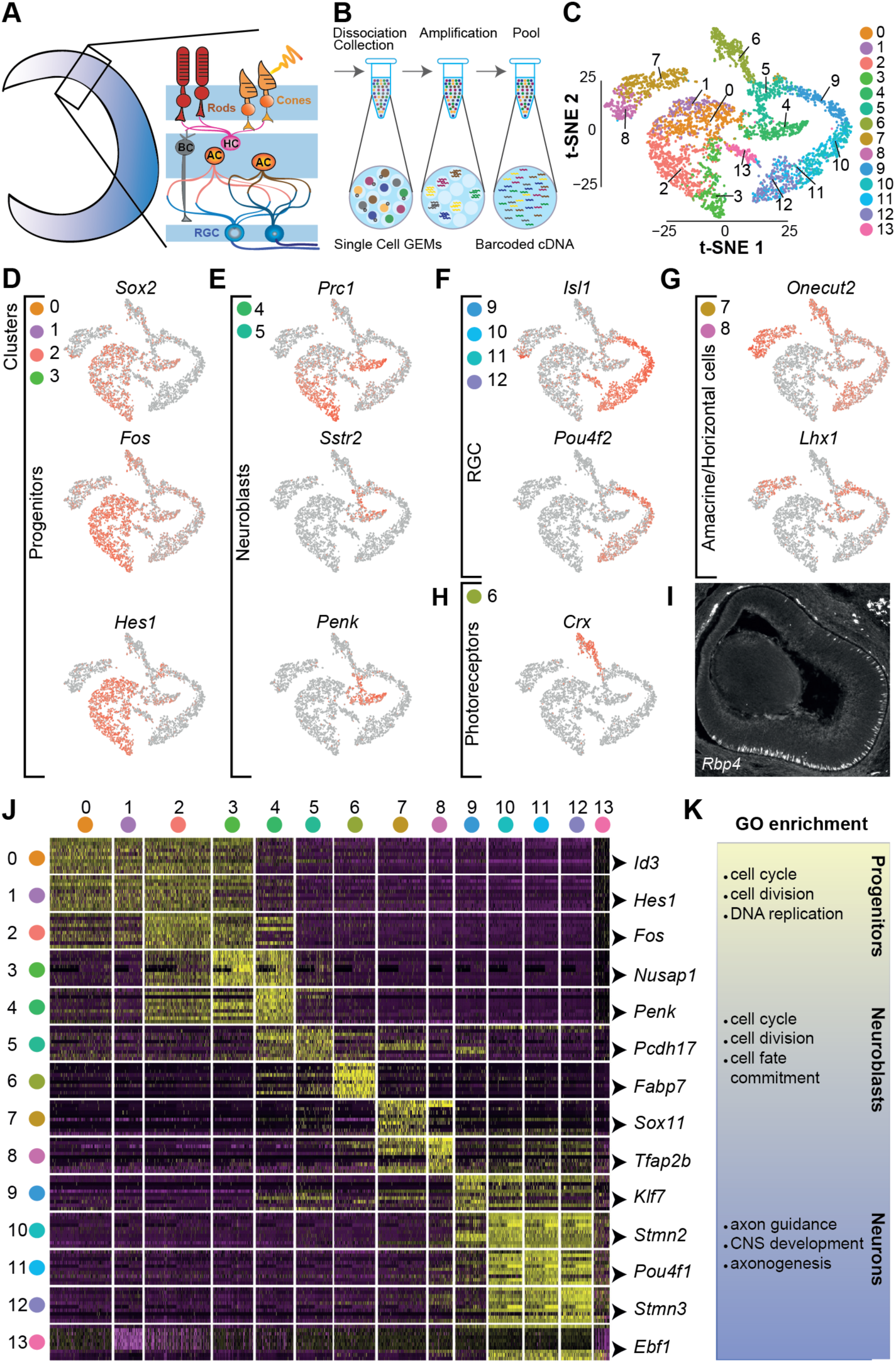
Developmental retina transcriptional diversity at the single-cell level. **A**. Schematic showing a developing retina, the layer organization of its cells and their diversity. **B**. Main steps of the droplet-based scRNA-seq 10X procedure. **C**. t-SNE reduction space of the 5348 cells transcriptomic profiles from the E15.5 retinas colored by the unsupervised clustering categories. **D-H**. List of markers genes colored by grey to red gradient representing the gene expression levels on the t-SNE that were used for the identification of the main cell-type clusters **I**. Signals of the Cre-reporter Tomato in coronal section detected from a *Rbp4-Cre; Ai14* retina at E15.5 with the layer of cones in the bottom part of the retina. **J**. Hierarchically organized heatmap of the top 10 most expressed marker genes for each cluster. **K**. GO terms associated for the main groups (progenitors, neuroblasts and neurons).

### Lineage reconstruction exposes hierarchical production of early neural fates

The global organization of the retinal clusters, as represented in **Fig. 1C**, was colored with the cluster identified by cell type, as identified in **Fig. 1**, with the RGC groups split in early, mid1, mid2 and late RGC based on their expression profile, resembling a lineage tree where most neuronal fates emerge from a neuroblastic state (**Fig. 2A**). In the retina, the generation of the neural fates follows a gradual shift of competence (Boije et al., 2014). In an attempt to better define the relationships between this peculiar cluster configuration, we first represented the transcriptional diversity using a uniform manifold approximation and projection (UMAP) as it facilitates the reconnection of divergent clusters (**Fig. 2B**) (McInnes et al., 2018). We were again able to observe a similar organization in which we saw that the progenitor pool is organized by cell cycle phase (**Fig. S4**). In this representation, the cluster assigned as AC and HC became connected to the neuroblast pool of cells that was also bridging to the RGC and cone photoreceptor clusters. Since the neuronal cells originated from the progenitor branch, we hypothesized that the extensions of these clusters were associated with their birthdate. As it was previously shown that the progenitor-to-neuron differentiation is organized along a pseudo-time axis with incremental steps in the three days following cell cycle exit (Telley et al., 2016), we then asked whether the extension of the clusters observed in the t-SNE and the UMAP would match their order along a pseudo-time axis (**Fig. 2C**). We observed a clear progression from the pool of progenitors (RPC) toward each of the clusters, with the first branch (purple) giving rise to the cones, the second branch to the AC/HC and finally the most mature group of cells at the extremity of the pseudo-temporal axis, the RGCs. We investigated whether the branches of this time-dependent organization would match a particular fate orientation (**Fig. 2D**). This was done by inferring the lineage sequence using a reconstruction of linear and branched trajectories based on Monocle 2 (**Fig. 2E**). While we saw a clear early commitment of cells toward the photoreceptors (most likely cones), the AC and HC, on a latter branch, were found to follow the same trajectory, suggesting a common progenitor. Further down the tree, the RGC segregated in two groups at N5 node, one from a pool of neuroblast expressing *Stmn1* and *Sncg* (right branch), and one mostly containing cells that we attributed to the CB cluster with many mitochondrial genes (left branch).

**Figure 2.**
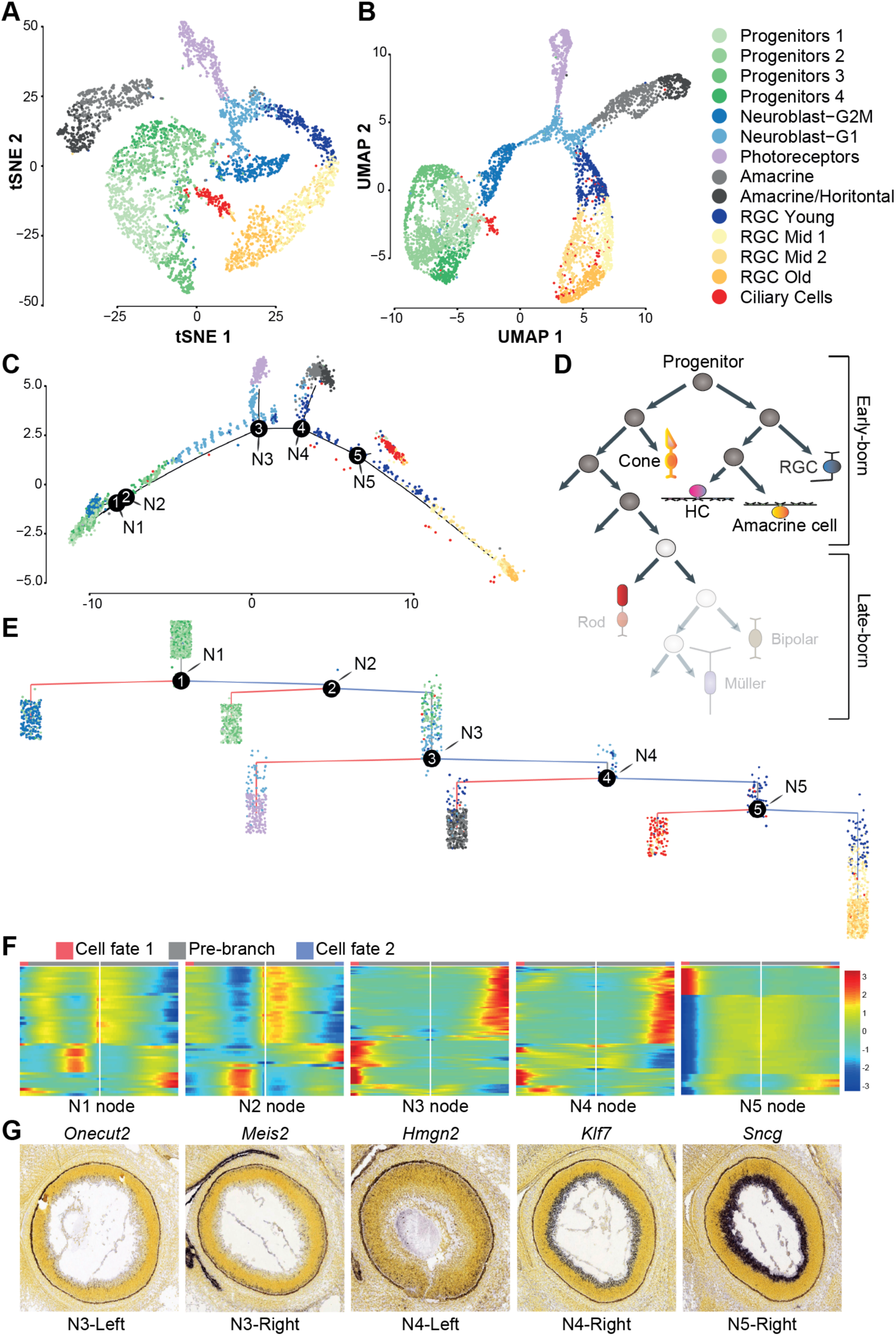
Early-born retinal neuron cell fate specification. **A**. Color-coded t-SNE of the 5348 cells based on cell type identification. **B**. Uniform Manifold Approximation and Projection (UMAP) dimension reduction of all cells. **C**. Pseudotime trajectory representation (monocle 2) showing five nodes at the root of seven terminal branches color-coded by cell type. **D**. Schematic of retinal lineages organization with the distinction between early- and late-born cells. The part of the lineage in transparency contain the late-born cells. **E**. Complex pseudotime lineage tree revealing the hierarchy between the five node points. N1 and N2 represents the progenitor’s nodes while N3 to N5 are the nodes giving rise to committed neurons. **F**. Heatmaps derived from Branched Expression Analysis Modeling (BEAM) show the dynamics of gene expression associated with fate orientation for the red-left branches (cell fate1) versus the blue-right branches (cell fate 2) of each node. Genes represented show significant differential expression values (q-value < 1.0e-20 in the BEAM test). **G**. ISH of influential genes in the determination of the associated neuronal node branching points.

In order to link cells from a progenitor pool to their corresponding fate we used the Branched Expression Analysis Modeling (BEAM) approach from the Monocle 2 pseudo-time analysis to trace back the gene sets responsible for the temporal transitions along the lineage tree groups (Hanchate et al., 2015; Qiu et al., 2017b). We plotted in heatmaps the dynamics of expression for the genes identified by BEAM to be the most prone to contribute to the balance toward one fate or another (**Fig. 2F** and **Fig. S6**). For the node N3, at the root of the cone group, we found the presence of *Casz1, Thrb* and *Meis2* to be among the top cone-oriented transcripts, while *Stmn2* and *Klf7* were scored toward the RGC/AC/HC fates. Further down the tree, the N4 node showed a misbalance with *Onecut2, Casz1* and *Ldhb* shifted to AC/HC group whereas *Stmn3* and *Elavl4* were two early predictors of the RGC/CB fate. Interestingly, we also revealed unbalanced expression of two miRNAs, *miR124a-1hg* and *miR124–2hg*, which preferential expression may trigger AC/HC fates (**Fig. S7**). Finally, at the last switch, we found mitochondrial genes for the CB branch versus *Ptma/H3f3a/H3fb* and *Sncg* in the RGC enriched branch. Interestingly, in the last segment a nonnegligible number of RGCs was also present in the CB group, suggesting a different transcriptional program for this sub-cluster of RGCs. We validated the expression pattern of some of the main influential transcripts by ISH (**Fig. 2G**).

Importantly, the node N3 also represents the junction from the progenitor pool to one of the four neuronal cell types, and this transition appears to emerge from a common root. Since this convergence is accompanied by a reduction of heterogeneity in the tSNE and the UMAP (**Fig. 2A-B**), we further explored the transcriptional dynamic of each cells by using RNA-velocity information (La Manno et al., 2018). Based on the ratio between nascent (unspliced) and mature (spliced) mRNA, RNA-velocity allowed us to tag each cell by a velocity vector (arrows) corresponding to its putative near future transcriptional state, further validating the progenitor-to-neuron organization (**Fig. 3 and Fig. S5)**. The transcriptional dynamics are particularly decreased at the intersection prior to neuronal cell type specification (**Fig. 3A**). Low velocities were also observed at the root of the progenitor cluster, then get robust directional flow towards each of the three neuronal branches to finally slow down again at the extremities (ends) of the neuronal cluster (**Fig. 3B**). We validated for early expressed genes the velocities (blue to red coloration on the UMAP) that predict the dynamics of gene expression in cells on their way for specification (**Fig. 3C-G**). *Dcc, Cplx2* and *Tfap2d* depicted pre-mRNA as early as the late progenitor cells while other transcripts such as *Stmn3* and *Tfap2b* started to initiate their production at the beginning of the RGC and AC branch, respectively. Altogether, these results reflect a sequential order in the production of the early-born retinal neurons, with the RGCs being in the most mature state of differentiation.

**Figure 3.**
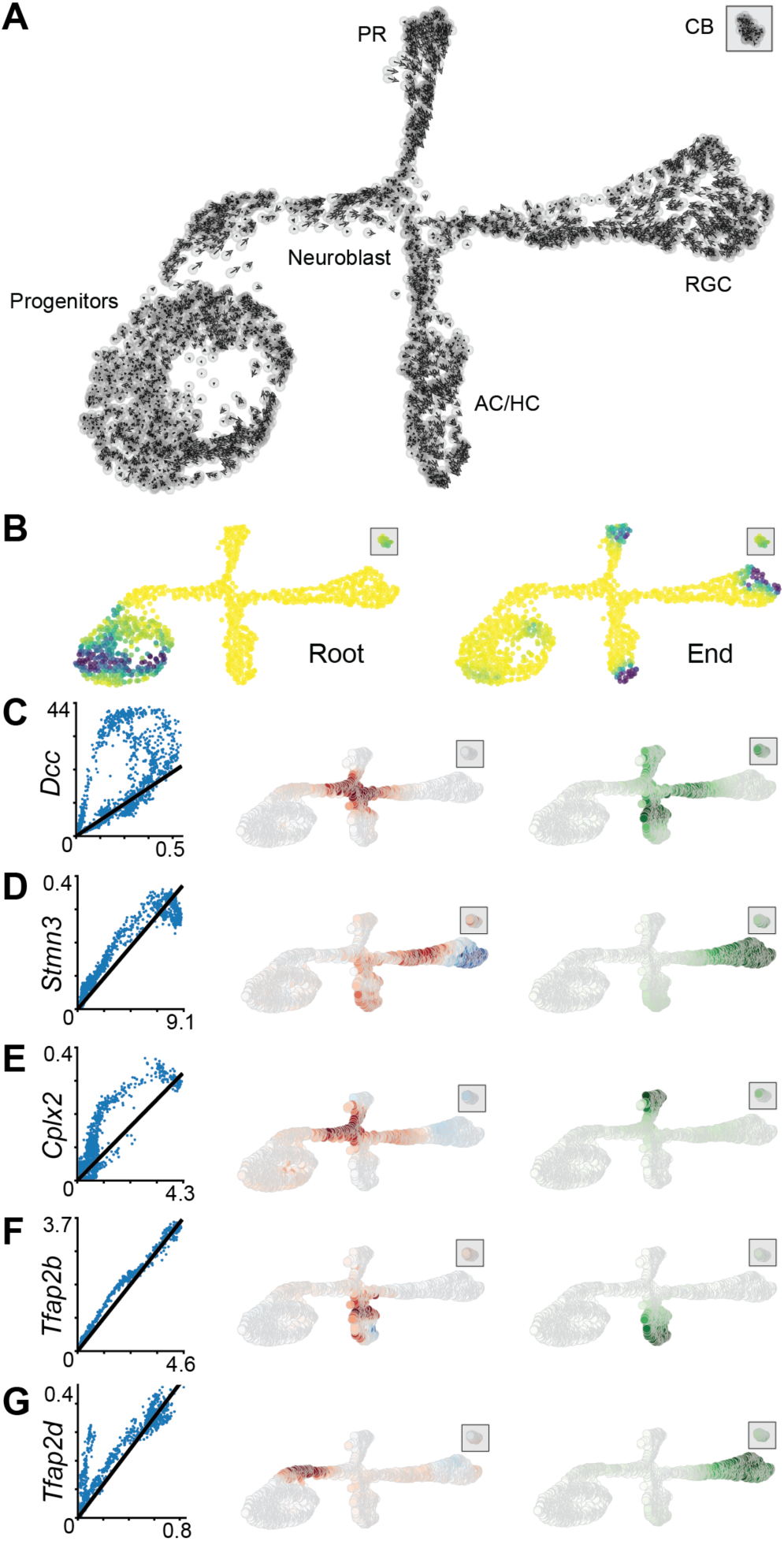
RNA velocity reveals directional progression of transcriptional states across retinal single-cells. **A**. RNA - velocity plot embedding in the UMAP space showing the putative near transcription state of each cell. **B**. UMAP color - coded by root and end points of the velocity representation, representing the starting points and the end points of the dataset differentiation process. **C - G**. Left curves are the phase portraits that show the variation of the ratio between unspliced (Y axis) and spliced (X axis) levels of mRNA for the pan neuronal marker (*Dcc*), as well as genes specific for RGC (*Stmn3* and *Tfap2d*), cones (*Cplx2*) and AC/HC (*Tfap2b*). Central plots are the corresponding velocity variances that are represented on the UMAP by color - coding from blue (decrease) to red (increase) of the gene expressions. The UMAP on the right show the level of gene expression in the single cells from white for absence to dark green for highest expression.

### Transcriptional waves drive retinal ganglion cell differentiation

Subsequently, we used the RGC clusters to study in depth the transcriptional programs at work during early neuronal differentiation. In order to sub-classify this large group of 1312 cells identified as RGCs, we first performed a principal component analysis (PCA) only on the RGCs to identify the main genes responsible for their transcriptional heterogeneity (**Fig. 4A**). We found that the principal GO categories segregating the RGCs were “cell differentiation” (PC1), “nervous system development and transcription regulation” (PC2) and “cell differentiation and survival” (PC3). The PC1 was responsible for more than 7% of the variance with a seeming temporal progression allowing compartmentalizing the RGCs based on their differentiation level, that is, reflecting their birthdate within the retina. This parameter gave us the possibility to cluster patterns of gene expression along the temporal progression. Genes presenting noticeable expression variations along the RGC were regrouped as time-dependent waves of transcription (**Fig. 4B**). We identified six transcriptional waves from 383 genes that were dynamically expressed along the main axis and represent excellent candidates for the coordination of differentiation of RGCs, including six well-characterized neuronal markers that we plotted on a UMAP of the pool of RGC only (**Fig. 4C-D, Table S2**): *Dlx1, Pou4f1, Tshz2, Pou6f2, Dnm3* and *Eomes*. Fifty-eight of these genes are transcription factors that are highly inter-connected in gene networks (**Fig. 4E**). Interestingly, we found they segregate in three groups. The first group consists of genes found enriched in cell cycle exit, with *Fos/Jun* and *Notch1/2* in its core (**Fig. 4E**, top, green circle), and was mostly connected to a second group of developmental transcription factors with *Neurod1* and *Isl1* as main interactors that were previously shown to specify RGCs (middle cluster, pink). This middle group of genes was slightly connected through *Pou4f2* (**Fig. 4E**) to a more distant cluster of genes that pattern synaptic terminals (e.g. *Snap25*), thus representing an advanced level of maturation. Taking *Pou4f2* and *Eomes*, two genes from the start and final waves, respectively, we observed a peripheric- and neuroblast layer-specific pattern for *Pou4f2* while *Eomes* was indeed observed in a more central position, a feature of mature RGCs (**Fig. 4F**). While the transcriptional waves allowed us to identify general differentiation programs, we tried to see whether we could identify micro-clusters of cells based on the expression of genes known to be expressed in single RGC subtypes. Among these genes, we found *Pcdh9* (Martersteck et al., 2017), *Slc6a4* (*Koch et al., 2011*) and *Prdm16* (*Groman-Lupa et al., 2017*). Their expression may be more restricted later on during RGC development (*Pcdh9*), or expand to other RGC cell types (*Slc6a4;* (Assali et al., 2017)). This latter gene is specific to the RGCs that will send their axon to the ipsilateral side of the brain (Koch et al., 2011; Peng et al., 2018). Overall, this analysis showed us that, at this stage (E15), we can recover a large spectrum of differentiation stages of an early retinal cell type.

**Figure 4.**
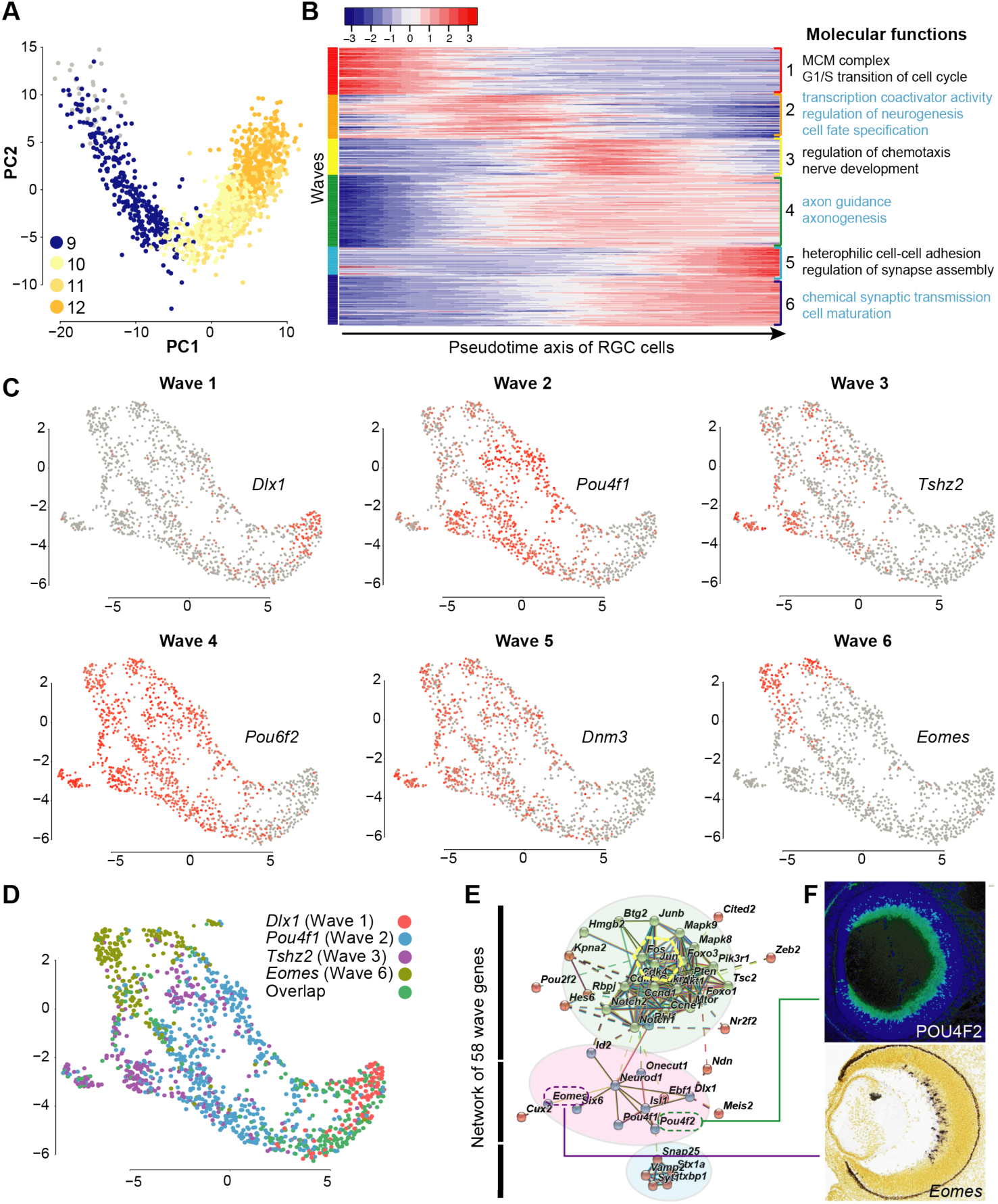
Retinal ganglion cell differentiation program follows transcriptional waves. **A**. Principal component analysis (PCA) on the 1312 RGC cells colored by clustering categories. **B**. Transcriptional waves patterns of RGC differentiation through pseudotime with on the right their associated GO term enrichment (molecular functions, derived from DAVID). **C**. UMAP colored for representative genes expression identified in the RGC wave analysis. **D**. Summarizing color-coded UMAP for representative genes of the cell groups from the four sharp waves (1, 2, 3 and 6). **E**. Network representation showing three module of gene organization based on their role and interactions in the wave dynamics. **F**. ISH and immunofluorescence that validates the rather peripheric/progenitor-pattern for POU4F2 (up wave 1) and the central position of *Eomes* (down wave 6) in E15.5 retinas.

### Central-to-periphery gradients unveil axon guidance ligand-receptor pairs

As retinal neurogenesis follows a central-to-peripheral gradient, we reasoned that molecules that are surface-bound or secreted in higher amounts by the more mature RGCs located in the central retina would be the best candidates to attract the axons of the more recently produced RGCs that are located at the periphery (**Fig. 5A**). Conversely, cues produced in higher amounts by young RGCs may play a repulsive role in an autocrine manner for their axons fleeing toward the optic nerve entry in the central part of the retina. In order to classify the central-to-peripheral position of RGC cells we took advantage of our differentiation analysis to attribute a “birthday” status to each of them. We identified eleven ligand-receptor pairs (**Fig. 5B**), with two of them (*Slit2/Robo2*, and *Dcc/Netrin*) being known to play a role in these guidance steps (**Fig. 5B-C, Table S4**) (Deiner et al., 1997; Thompson et al., 2009). We listed the top segregating genes and plotted them along the maturation axis in partitioned UMAPs of RGCs (**Fig. 5C**). Among the ‘old’ genes we found a positive control, *Igf1*, which is a secreted ligand that was shown to be strongly expressed in the more central part of the retina (Wang et al., 2016). On the other hand, ‘young’ genes included *Igfbp5, Igfbpl1, Dlx1* (**Fig. 5C, Table S3–4**). Among the other secreted ligands, *Reelin* mRNA was expressed in a gadual manner from periphery to center, so did *Kitl* transcripts (**Fig. 5B, D**). Of note was also the strong detection in RGCs of *Tenm3*, a transcript important for ipsilateral RGCs, as well as *EphB1*, a gene coding for a guidance receptor involved in the segregation of ipsilateral RGCs at the optic chiasm (**Fig. 5E**) (Leamey et al., 2007; Williams et al., 2003). Beyond the molecules known to play a direct role in axon guidance, we also identified genes encoding for putative downstream machinery, including the central-high gene *Pde4d* and the periphery-enriched gene *Btg2* (**Fig. 5F-G, Table S3–4**). We found complementary patterns for *Slit2/Robo2* and the Netrin-1 receptors (*DCC/Unc5*), which we plotted in the same tSNE (**Fig. 5H-I**). Together, these results validate previously known genes and identify new ones with expression pattern that suggest a role in retinal axon guidance.

**Figure 5.**
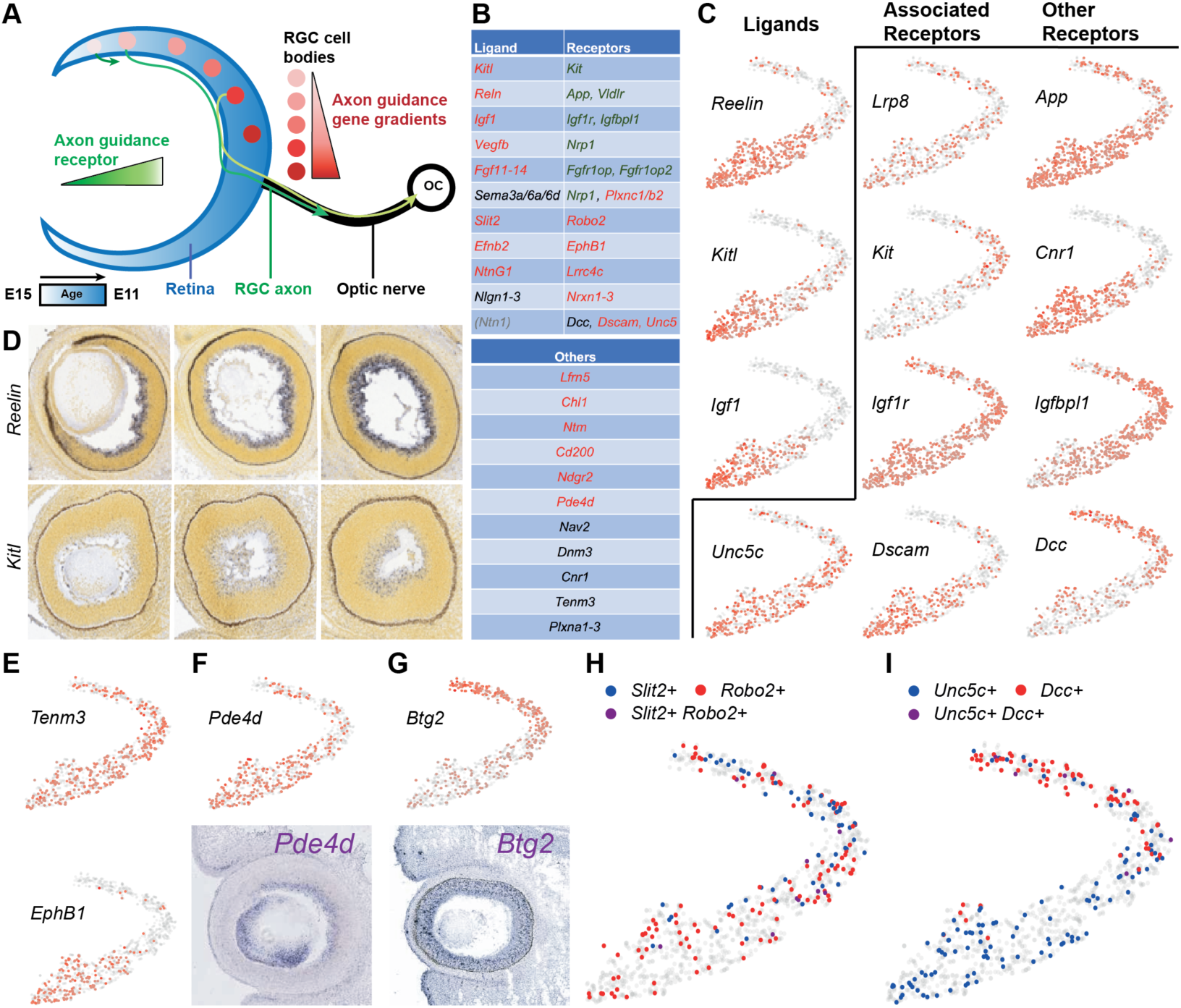
Central-to-periphery gradients unveil axon guidance ligand-receptor pairs. **A**. Schematic explaining the axon guidance cues gradients detection hypothesis. **B**. List of important axon guidance genes including eleven pairs of ligand-receptors identified in RGCs. **C**. Colored t-SNE from RGCs showing matching pairs. **D**. ISH of *Reelin* (up) and *Kitl* (down), two of the main ligands distributed in high central-low-peripheric gradients. **E**. Colored t-SNE for *EphB1* and *Tenm3*, two ipsilateral-RGC related genes. **F-G**. Color-coded t-SNE and ISH for putative novel expression axon guidance player: *Pde4d* (**F**, enriched in mature RGCs) and *Btg2* (**G**, enrich in early RGC). **H**. Color-coded expression levels of *Slit2* and *Robo2* in the same t-SNE showing their complementary patterns. **I**. Color-coded expression levels of the complementary patterns of the three Netrin-1 receptors in the same t-SNE showing *Dcc* in ‘young’ RGCs and *Unc5c* and *Dscam* in ‘old’ RGCs.

### Spatial reconstruction using retinal ganglion cell transcriptomes

Another prominent spatial feature of RGC is their position along the dorso-ventral (DV) and temporo-nasal axis (TN). We used this feature to address our second aim, the deduction of spatial information from transcriptional signatures. Importantly, the position of RGC in one of the four quadrants has been associated with their projection in the developing brain (McLaughlin and O’Leary, 2005). This concordance between the cell body position and their thalamic and colliculus synaptic targets being referred to as retinotopy and requires specific transcriptional programs. Here we exploited this particular feature to position the cells in a pseudo-spatial orientation. To do so, we first plotted the expression levels of known marker genes for the four different orientations (**Table S5)** (Behesti et al., 2006, 4; McLaughlin and O’Leary, 2005; Takahashi, 2003) on our RGCs that we segregated in five t-SNE dimensions (**Fig. S8** – DVTN t-SNE). We took advantage of these micro-clusters of cells to extract their transcriptional signature. E15.5 being the peak stage for ipsilateral axon segregation (Colello and Guillery, 1990; Drager, 1985; Guillery et al., 1995), we reasoned that RGCs projecting contralaterally and ipsilaterally would express different sets of axon guidance related-genes. We used two approaches. First, we extracted the cells with high ventro-temporal (VT) scores from the pseudo-spatial information (**Fig. 6**). Second, using an ipsilateral specific gene, *Slc6a4*, we analyzed the distinct subpopulations without any spatial considerations (**Fig. 7**). With the first approach, we found that the best spatial discrimination is observed between the first and the third dimension (**Figs. 6A-C and S8**). We subsequently attributed a VT, DN, DT or VN score to every cell that we plotted either in a t-SNE (**Fig. 6C**) or in two ordinal axes (**Fig. 6D**). This classification allowed us to perform a comparison between groups of transcriptomes from the four distinct quadrants. We identified 210 single cells that were attributed to the VT quadrant of the retina. Nineteen genes were differentially expressed between VT and other RGCs cells: *Crabp1, Nefl, Gal, Eno1, Cbx1, Rassf4, Ppp1r1a, Ssu72, Acp1, Rps12-ps3, Mid1ip1, Chchd2, Ckb, Bbx, Mdk, Prdx1, Ptprg, Gria1, Mfng* and *Btg2* (**Fig. 6E**). Finally, we performed a comparison between all the dorsal and ventral RGCs. In this analysis we found 1195 genes differentially expressed (**Table S6**) from which three were validated by ISH: two dorsal genes (*Cnr1* and *Rprm*) and one ventral gene (*Irx2*) (**Fig. 6F**).

**Figure 6.**
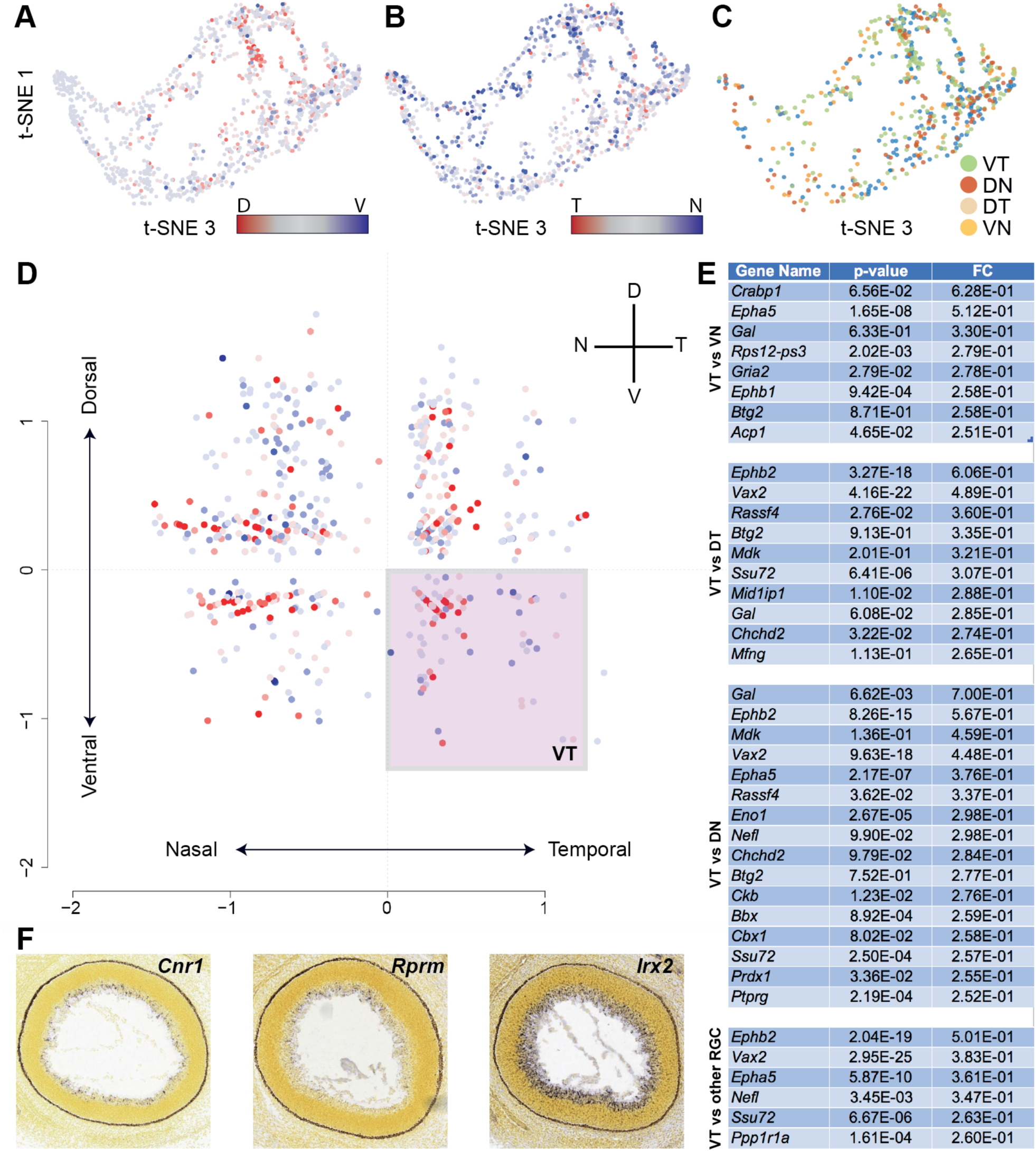
Spatial reconstruction using retinal ganglion cell transcriptomes. **A**. Dorso-ventral (DV) gene expression ratio (Q80) in a t-SNE from RGCs with X-axis t-SNE3 and Y-axis t-SNE1. **B**. Temporo-nasal (TN) gene expression ratio (Q80) in a t-SNE with X-axis t-SNE3 and Y-axis t-SNE1. **C**. Classification of cells based on their DV and TN scores in either four retinal quadrants (VT, VN, DT or DN). **D**. Retinal pseudo-space organization showing the fraction of VT cells in the box, and the four groups that were used for the gene expression analysis. **E**. List of genes differentially expressed between VT and the other quadrants. **F**. ISH of the newly identified dorsal genes (*Cnr1* and *Rprm*) and ventral gene (*Irx2*) in E15.5 mouse retina.

**Figure 7.**
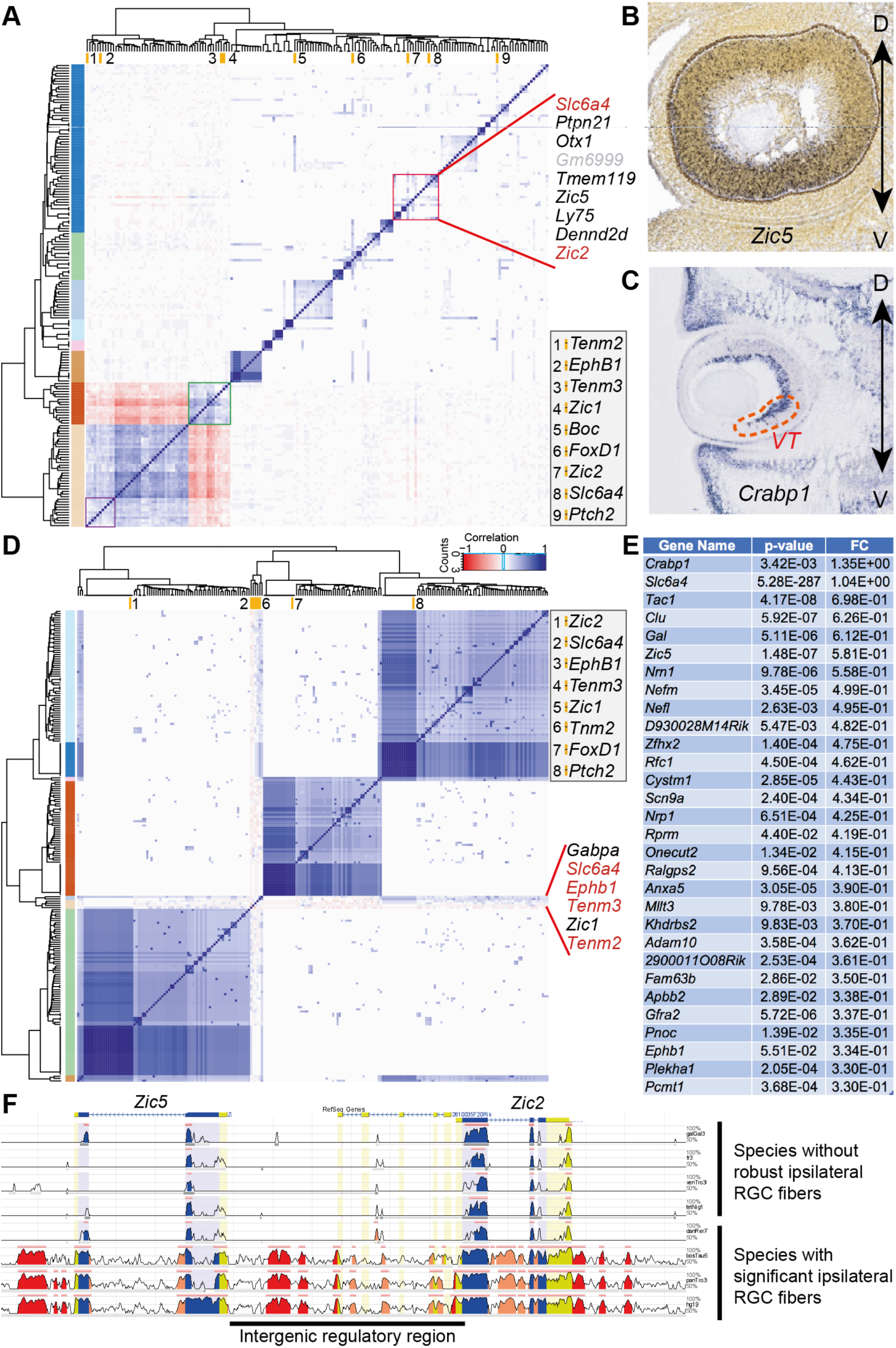
Transcriptional signature of ipsilateral-projecting RGC. **A**. Heatmap of co-expression from 218 genes co-expressed with ipsilateral genes in the 1312 RGCs. **B-C**. ISH validating the VT localization of *Zic5* (E13.5) and the ventral enrichment of *Crabp*1 mRNA (E14.5). **D**. Heatmap of co-expression from 99 genes co-expressed with ipsilateral genes in the 147 VT cells. **E** List of top 30 genes differentially expressed in the *Slc6a4* positive RGCs (28 cells) versus the other RGCs originating from the same RGC maturation window (182 cells). **F**. Regulatory landscapes of the newly identified candidate transcription factor for ipsilateral RGCs *Zic5* and its paralog neighbor *Zic2* (extracted from rvista 2.0).

To understand how the VT genes would be linked to previously known ipsilaterally-associated genes, we conducted a co-expression analysis to show the extent to which they are co-expressed in single-cells. Among the ipsilateral RGC genes, 8 were expressed in enough RGC cells (> 0.5% RGCs) to be included in the analysis, consisting of *Slc6a4, EphB1, Tenm2, Tenm3, Zic1, Zic2, FoxD1 and Ptch2* (Clark et al., 2018) (**Fig. 7A**). The main cluster comprises three of these genes, that were co-expressed with *Zic5* and *Crabp1*. By ISH we observed that *Zic5* had indeed ventral high expression and that *Crabp1* had even VT high expression, thus strengthening a potential role in ipsilateral RGC identity (**Fig. 7B-C**). However, we noticed that these known ipsilateral genes were not frequently co-expressed in the 1312 RGCs (**Fig. 7A** and **Fig. S9**), a feature that was also observed in E14 and E16 time points from similar datasets **(Fig. S9**) (Clark et al., 2018). When the analysis was restricted to the VT RGCs (92 cells), we found that five of these 8 genes (*Slc6a4, EphB1, Tenm2, Zic1* and *Tenm3*) were tightly associated in a hierarchical clustering organization, in a small and central cluster (**Fig. 7D**). In this analysis we found that the transcription factor *Gabpa* was strongly associated with this group. Finally, in the RGCs positive for *Slc6a4* (28 cells), we perform a differential expression analysis and found a slight enrichment for seven genes (**Fig. 7E**). Among them were *Zic5* and *Crabp1*, further validating our aforementioned co-expression analysis. Interestingly, the regulatory landscape near to *Zic5* highlights common regulatory elements with the ipsilateral RGC master gene *Zic2* **(Fig. 7F)**. Altogether, these results show that RGC position can be inferred quite confidently from their single-cell transcriptomes and that a small subtype such as the ipsilateral RGC can be identified.

## DISCUSSION

Here we provide a resource of 5348 retinal single-cell transcriptomes allowing deciphering the stepwise progression of gene expression during the generation of early-born retinal cells. In particular we used the RGCs as the most populated and robust group to delineate the logic of this progressive maturation. We discovered that transcriptionally inferred spatial information can be used to define subgroups of retinal cells that project differentially, leading to the identification of novel candidate genes playing a role in their extremely diverse arborization patterns.

### Early retinogenesis: pre-patterning versus stochasticity?

Our lineage analysis consolidated the existence of a hierarchy in the production of the early-born retinal cell types. Interestingly, when we search for genes that control the binary fate decision, we observed an early segregation of progenitors giving rise to the four classes of early-born neurons (i.e. downstream of Node1, **Fig. 2E**) versus the one giving rise to other cell types (branching on the left sides of N1 and N2). This aligns well with previous findings showing a shift in competence between the two phases of production, a phenomenon shown to be intrinsically driven (Belliveau and Cepko, 1999; Reh and Kljavin, 1989). The cone branch appears to be quite homogenous, in agreement with a recent study showing a production by symmetrical terminal divisions (Suzuki et al., 2013). In agreement with previous studies, we further emphasize a role for *Otx2*, *Neurod4*, *Casz1* and *Rpgrip1* in the generation of photoreceptors (Emerson et al., 2013, 2; Hameed et al., 2003; Mattar et al., 2015). The primary activation of *Otx2* and *Neurod4* in progenitors predicted well an early fate bias towards the photoreceptors (**Fig. S6** Node 3). The progenitors upstream of that branch (above N3) seem to have the competence to generate both cones and the other early-born cell types such as HC and RGCs. This latter observation supports previous lineage analysis from labelled clones in the chicken and mouse where a common progenitor can give rise to both HC and cones, while AC and HC seem to originate from a common transcriptional program (Boije et al., 2014). Finally, the progenitors giving rise to both RGC and potential CB cells also participate in the emergence of progenitors producing RGCs only. Under this node, cells segregated in two groups, one from a pool of progenitors expressing *Stmn1* and *Ptma* (the RGCs only branch), and the other one mostly containing cells expressing high levels of mitochondrial genes that were attributed to the CB cluster. This finding is in agreement with a previous study which showed, using lineage tracing or time-lapse microscopy in transgenic mice, that a subset of RGCs originated from the ciliary margin zone (Marcucci et al., 2016; Wetts et al., 1989).

### Chromatin remodeling and metabolic rate to pave the way for cell fate specification

Relaxing chromatin prior to transcriptional activation is a balance between pioneer transcription factors and the action of chromatin remodelers. Our study, with its well-defined time dimension from progenitors to the early phase of cell fate specification, is a key resource for the identification of these upstream actors of neuronal differentiation. Here, we showed that early genes that are dynamically expressed during the first phase of RGC differentiation include many transcription factors and chromatin remodelers. Among them *Hdac, Ktm2, Ezh2, Hmga1/2* (AC), *Hmgb2*, and several *Prdm* transcripts are known to be major actors in transcription regulation (**Fig. S6, S10, Table S3–4**). Their expression pattern precedes the remodeling of chromatin and may serve as a permissive signal to restrict multipotency and cell fate commitment (Chen and Cepko, 2007; Fabre et al., 2018; Iida et al., 2015, 2; Pereira et al., 2010, 2). Among the *Prdm* gene family, a family related to the catalytic SET methyltransferase domain, while *Prdm1* and *Prdm13* are found specific to cones and AC cluster (respectively), *Prdm2* and *Prdm10* dynamically define RGC subpopulations, a feature known for *Prdm16* (Groman-Lupa et al., 2017). Intriguingly, our analysis also shows enrichment for metabolic genes. For example, while the lactate dehydrogenase gene *Ldhb* is enriched in the AC/HC and RGC clusters, *Ldha*, which is at basal levels throughout the dataset, is more strongly expressed in the cone photoreceptors emerging cells. It may highlight a function for aerobic glycolysis process in cone photoreceptors to meet their high anabolic needs even at the beginning of their maturation (Chinchore et al.; Zheng et al., 2016) (**Table S1, Fig. S6, S11**). In the last node (N5), the binary choice is associated with a misbalance of mitochondrial transcripts (**Fig. 2E**, **S6**). This asymmetric distribution might be a consequence of an increased number of mitochondria and thus reflect a differential mode of metabolism in this cluster. A similar process with asymmetric distribution of aged mitochondria have been shown to influence the stemness or stemlike cells (Katajisto et al., 2015). How this differential expression influences RGC fate remains to be established.

To follow the establishment of cell fate specification, it is essential to delineate the chronological orders in which transcription factors are produced, that is, to understand their hierarchical organization, and thus reconstruct a clear gene regulatory network. In this study we took advantage of a recently developed procedure to unveil forms of pre-mRNA produced by a cell giving a direction for each cells representing a prediction modellable on the UMAP space (**Fig. 3 and S5**) (La Manno et al., 2018). We observed that neuroblasts depict a transcriptional bottleneck profile: their velocity vectors are decreased near the main branching area of the UMAP space, implying a common loss of gene expression dynamics followed by a secondary multi-directional diversification, as it was shown for oligodendrocyte precursor cells in neocortex (Zeisel et al., 2018). Using this framework, we established a more precise chronological sequence between the transcription factors identified along the pseudo-time axis, giving further strength to our gene network.

### Spatial information as a proxy for RGC axon guidance transcriptional programs

Our results led us to postulate that the early-born neurons that project in an untracked environment (i.e. they cannot rely on fasciculation-like mechanisms) are equipped with a peculiar set of guidance receptors in order to navigate the CNS. One well-characterized neuronal cell type that is produced early, with diverse and long-range projections, are the RGCs (Petros et al., 2008; Sanes and Masland, 2015; Seabrook et al., 2017; Trimarchi et al., 2008). We hypothesized that RGCs with an older ‘birthday’ status (at the extremity of the pseudo-time axis) would be the ones born between E11 and E12 and would thus already be making connections with distant brain territories such as the dorsal lateral geniculate nucleus and the superior colliculus, two regions where they are known to segregate based on their location in the retina (Seabrook et al., 2017).

RGC projecting to different brain territories originate from different quadrants of the retina, a phenomenon known as retinotopy. We thus took advantage of our DV and TN classification to compare their transcriptional signatures (**Fig. 6D-E**). Previous studies have shown that the retinotopy of the retina is associated with differential expression of *Ephrin* family members (Cang et al., 2008; McLaughlin and O’Leary, 2005). Here we validated these groups of cells, including *Crabp1* that was found to be highly enriched in the ventral segment of the retina (Díaz et al., 2003). We further show new molecular players acting with these membrane-bound guidance receptors.

Our study led us to identify a dozen of ligand and receptors, including a few for which the functional roles had been previously validated in mutant retina (**Fig. 5B**). For instance, the peripheral-*Slit2*/central-*Robo2* complementary expression pattern, a pattern that was already detectable by ISH (Erskine et al., 2000), was implicated in the fine wiring of the RGC axons on their path toward the optic nerve, with a requirement exclusively for RGC axon guidance within the peripheral retina (Erskine et al., 2000; Thompson et al., 2006). This phenotype seems now compatible with a lack of repulsion from the periphery to the middle part of the retina, where attractive cues may take the relay to stir the axons. Two of the ligand-receptor systems we identified are the *Kitl/Kit* pairs and the *Netrin-1/Dcc* signaling (**Fig. 5B-D**). The former pair encodes for SCF and its receptor and was never shown to be involved in guiding retinal neurons, thus further functional characterization will be required to confirm this effect (Williams et al., 1990). However, their function as axon outgrowth-promoting cues was previously established in the peripheral nervous system (Gore et al., 2008; Hirata et al., 1993). In the retina, as the expression of the ligand (*Kitl*) increases along the RGC maturation axis (**Fig. 5C**), the expression of its receptor (*Kit*) is mostly detected in younger RGCs. Interestingly, the combined and gradual expression of *Adam10* (**Fig. S12**), that encodes a protease processing SCF in its soluble form, from the early to the late RGCs, may participate in the establishment of a gradient of the ligand along the central-to-periphery domains. For the latter pair (*Ntn1-DCC*), we did not observe the *Ntn1*-expressing cells but we clearly detected *DCC* as strongly expressed in ‘young’ RGCs, at a time they most likely send their axons toward the center of the retina where the source of netrin has been documented (**Fig. 5C**) (Deiner et al., 1997). While the outgrowth-promoting activity of Netrin-1 was shown to be abolished by DCC-blocking antibody (Deiner et al., 1997), it was unclear then how the axons would escape this netrin attraction to pursue their route in the optic nerve. As axons can have opposite turning responses to Netrin-1 depending on the status of cytosolic cAMP-dependent activity (Ming et al., 1997), the switch in Netrin-1 activity was then shown to rely on such mechanisms depending on the age of RGCs (Shewan et al., 2002). In our study, we identified two other Netrin receptors that may influence this switch. While *DCC* is expressed in young RGCs, *Unc5c* and *Dscam*, encoding for two receptors that were shown to heterodimerize to mediate growth cone collapse (Purohit et al., 2012) are switched on while RGCs get older, concomitant with the extension of their axons that have passed the optic nerve entry (**Fig. 5C, I**). How the switch from *DCC* to *Unc5c/Dscam* is orchestrated remains to be established.

### Identification of ipsilateral RGC signatures

Perhaps the smallest subpopulation of RGCs that we could identify was the ipsilateral RGCs. A recent study showed that after retrograde tracing to distinguish the RGCs that have crossed from those that have not crossed, the two populations depict different transcriptional signatures that remain after crossing the chiasm (Wang et al., 2016). Here, we used *Slc6a4* as an early marker of ipsilateral RGCs to identify the transcriptional set of mRNA that accumulated prior to the crossing. While we fail to detect *Zic2*, a transcript encoding a master transcription factor first identified for its patterning role in ipsilateral RGCs (Herrera et al., 2003), we did detect *Zic1* that was shown to be enriched in the retrogradelly labelled ipsilateral RGCs (Wang et al., 2016).

Moreover, in RGCs from the ventro-temporal segment, our analysis showed enrichment in specific factors involved in the maturation of RGCs with *Zic5* and *Crabp1. Zic5* was also closely co-expressed with *Slc6a4*. As this gene shares many regulatory elements with *Zic2* (**Fig. 7F**), it may also have similar targets and thus play a role in the specification of ipsilateral RGCs. Of note, we fail to detect other genes involved in ipsilateral RGC guidance, such as *Boc* (Fabre et al., 2010; Peng et al., 2018) and *FoxD1* (Carreres et al., 2011, 1; Herrera et al., 2004), in a sufficient number of cells. Along with *Zic2*, these genes might be either expressed at levels below detection, or be subject to high drop-out events, a caveat of the drop-seq procedure. Further study using more sensitive protocols may be useful to further identify an exhaustive list of genes specific to micro-clusters such as the ipsilateral RGC. The discovery of new genes that are important to establish and maintain axonal connections using single-cell RNA-seq data is becoming a promising avenue. The use of recent barcoding strategy to address it may further expand our knowledge in that sense (Klingler et al., 2018).

## Conclusions

In this study we have presented a fundamental resource of single-cell transcriptomes in the developing retina. We showed that retinal progenitors exhibit a high level of transcriptional heterogeneity and unveil its meaning with the identification of all the early born cell types. Moreover, we show that among these cell types we can resolve fine-scale diversity with, particularly for RGCs, the identification of differentiation waves, spatial position and meaningful gene matching important function such as the patterning of circuit formation.

We believe this study will facilitate our understanding of how the retina develops, especially in terms of cell fate specification for the early-born neurons. It paves the way for many functional perturbations related to sequential generation of early retinal cell types, the transcriptional dynamics leading to cell differentiation, and finally the temporo-spatial coordination of axon guidance programs. Finally, the refinement of gene regulatory networks and co-expression analysis has revealed deep relationships between retinal genes, and thus represent an important basis for a better understanding of retinal development at the single-cell level.

## Supporting information

## Acknowledgements

We thank Gioele La Manno for his help in the implementation of the RNA velocity. Nicolas Lonfat and members of the Jabaudon lab for critical reading of the manuscript, as well as Alexandre Dayer and Denis Jabaudon for their advices and sharing mice and reagents. We also thank Audrey Benoit, Christelle Borel and Wafae Adouan for assistance in the 10X procedure. P. Abe helped with the maintenance of the *Rbp4-Cre* mouse line, and N. Baumann for the generation of the 3D UMAP animation. We thank M. Docquier and D. Chollet and the Geneva Genomics Platform (University of Geneva).

## Authors contributions

P.J.F. designed the project. Q.L.G. and P.J.F. conceived and performed the experiments. Q.L.G., M.L. and P.J.F. performed the bioinformatic analysis. P.J.F wrote the manuscript with input from all authors.

## Data availability

The datasets generated and analyzed for this study are available in the GEO repository under accession number GSE12246.

## MATERIAL AND METHODS

### Mice

Experiments were performed using C57Bl/6 (Charles River), Ai14 Cre reporter (Jackson Laboratory #007914) and Rbp4-Cre mice that were bred on a C57Bl/6 background. Tg(Rbp4-Cre)KL100Gsat/Mmcd (denoted as Rbp4-Cre, GENSAT RP24–285K21). Embryonic day (E) 0.5 was established as the day of vaginal plug. All experimental procedures were performed at E15.5 and were approved by the Geneva Cantonal Veterinary Authority.

### Single-cells preparation

Coordinated pregnant mice of 12 to 30 weeks were ethically sacrificed to extract E15.5 embryos eyes. Thirty retinas were extracted in ice-cold L15, micro-dissected under a stereomicroscope and incubated in 200μl single cell dissociation solution consisting in a papain (1mg/ml) enriched HBSS at 37°C for 12 minutes with trituration every 2 minutes. At the end of the incubation time, cells were further dissociated via gentle up-and-down pipetting. Reaction was stopped with the addition of 400μl of ovalbumin enriched cold HBSS and the cell suspension was then passed on a 40μm cell strainer to remove cellular aggregates. Cells were then centrifuged 5 min at 500 G at 4°C. After removal of the liquid, the pellet was suspended in 250μl of cold HBSS and the resulting solution was finally FAC-sorted on a MoFloAstrios device (Beckman) to reach a concentration of 410 cells /μl.

### Droplet-Based scRNA-seq

The libraries of single-cells were prepared using the Chromium 3’ v2 platform following the manufacturer’s protocol (10X Genomics, Pleasanton, CA). Briefly, single cells were partitioned into Gel beads in EMulsion (GEMs) in the GemCode instrument followed by cell lysis and barcoded reverse transcription of RNA, amplification, shearing and 5’ adaptor and sample index attachment. On average, approximately 5,000 single cells were loaded on each channel with 2675 cells recovered in the first replicate (index F2), and 2673 cells recovered in the second (index E2). Libraries were sequenced on a HiSeq 4000 (Paired end reads: Read 1, 26bp, Read 2, 98bp).

### Importation and normalization

After the CellRanger (Version 2.1.1) processing on the latest mouse assembly (mm10), Seurat package version 2.3 (Butler et al., 2018) was used to import in R version 3.4.4 (R Core Team) 2 batches of 2673 and 2675 cells to perform downstream analysis. The merge of batches was performed by aggregation with the inbuilt Seurat function MergeSeurat while batch effect correction was made possible by linear regression of the transcriptomic expression of the 2 batches during the scaling and centering of the dataset by the ScaleData function.

### Filtering

Cells considered during the creation of the Seurat object expressed at least 200 genes, and genes kept are expressed in a minimum of 3 cells. Mitochondrial gene effect was regressed out for the whole data. 1648 variable gene were defined on variability plot as mean expression above 0.1 and dispersion above 1.

### Dimensionality reductions and cluster analysis

Principal component analysis (PCA) was then performed on these variable genes to reduce dimensionality of the dataset. Spectral t-distributed stochastic neighbor embedding (Spectral t-SNE) was based on the reduced dimensional space of the 5 most significant dimensions of the PCA using the Rtsne package Barnes-Hut implementation of t-SNE forked in Seurat with a perplexity set at 30. Dimensions used 1:5, default parameters. (Maaten, 2014). A t-SNE based clustering analysis was then performed by the shared nearest neighbor (SNN) modularity optimization algorithm (Maaten, 2014). Differentially expressed genes between clusters were obtained by Seurat implemented Wilcoxon rank sum tests with default parameters. The cluster identities in this t-SNE-space were uncovered by feature plots of typical cell types markers genes. Another approach of the dimensionality reduction problem was explored by Uniform Manifold Approximation and Projection (UMAP) (McInnes et al., 2018), generated with the help of Seurat and the UMAP-learn python package on the 10 most significant dimensions of the PCA. Three embedded dimensions of the UMAP were outputted for further use to represent the pattern of various features during differentiation. The minimal distance parameter of the UMAP was set to 0.3, number of neighboring points used in local approximations was set to 30 and the distance metric used to measure the distances in the input space was correlation. GO term analysis were performed using the DAVID bioinformatics resources 6.8. (Huang et al., 2008; Huang et al., 2009).

### Pseudo-temporal analysis

Pseudo-time analysis was performed using Monocle 2 using genes that have passed the QC of the Seurat object creation and E2 experiment cells (Qiu et al., 2017a). Genes considered as defining the progress of the pseudo-time were those that were detected as having an expression above 0.5 by Monocle. Negative binomial was considered for the model encoding the distribution that describes all genes. During the pseudo-time processing, the dimensionality of the dataset was reduced by the Discriminative Dimensionality Reduction with Trees (DDRTree) algorithm on the log-normalized dataset with 10 dimensions considered. Thus, branched expression analysis modeling was performed on major branching points with default parameters.

### RGC cluster analysis and waves analysis

The RGC cluster was analyzed (merging, normalization, batch correction, dimensionality reduction techniques and differentially expressed genes analysis) as previously described using Seurat with the exception of variable gene defined on variability plot as mean expression above 0 and dispersion above 0.8. The RGC waves analysis was performed with default parameters as described previously (Telley et al., 2016). Genes presenting interesting variations across RGCs were regrouped along a pseudotime axis, forming clusters composed of similar time-dependent gene expressions. Theses clusters of patterns were labeled as waves one to six.

### RNA velocity

Computation of RNA velocities uses the abundances of spliced and unspliced RNA to fit a model of gene expression and to estimate the rate of change of expressions in time across the whole transcriptome. This model is then used to extrapolate the short future gene expressions of each cells of the dataset. We used velocyto CLI to obtain separated counts for spliced and unspliced molecules and we used velocyto analysis module to calculate and visualize velocities following the same filtering and pre-processing procedure as done in the original study (La Manno et al., 2018). We processed the reads of the single-cell experiment with the run10x function on the loom file outputted from the CellRanger Pipeline, with reference genome of the mouse, mm10 (Ensembl 84), from 10x Genomics, and masked the corresponding expressed repetitive elements that could constitute a cofounding factor in the downstream analysis with the UCSC genome browser mm10_rmsk.gtf. Downstream analysis and representation of velocities were performed by two packages for validation: the scvelo standard pipeline for the supplementary figures and the Velocyto python packages for main figures (La Manno et al., 2018). We used the Jupyter notebook DentateGyrus.ipynb as a guideline for the latter with default parameters for fitting gene models, normalize and represent the data with following exceptions: The 15 first principal components of the PCA were used in analysis and the matrix were smoothed by k-nearest neighbours (knn) using 180 neighbours. RNA velocity transcriptional dynamic was embedded on a UMAP space computed with the UMAP python package, parameters set for a number of neighbours of 120, a learning rate of 0.5 and a min distance of 0.4.

### DVTN scores

For each RGC cell, a Dorso-Ventral (DV) and a Temporal-Nasal (TN) score was calculated as the log2 ratio of the quantile 80 of the Dorsal genes (resp. Temporal genes), with the quantile 80 of the Ventral genes (resp. Nasal). A pseudo-expression of 1 was used to avoid infinite values. Those scores were used to classified the RGC cells into 4 groups: “DT” (DV > 0 & TN > 0), “VT” (DV < 0 & TN > 0), “VN” (DV < 0 & TN < 0), and “DN” (DV > 0 & TN < 0). Only cells with a non-null DV or TN score were classified (606 cells). Differentially expressed genes between the four cell groups were obtained by Seurat implemented Wilcoxon rank sum tests with default parameters.

### Co-expression analysis

The heatmap in **Figure 6** represents the genes with an absolute Spearman’s correlation greater than 0.2 with at least one of the 9 RGC-ipsi genes (176 genes). Only cells expressing one of those RGC-ipsi genes were considered for the correlation analysis (781 cells). The 8 clusters shown on the heatmap have been identified with a complete-linkage clustering based on an euclidian distance.

### ISH and IHC (Immunohistochemistry)

Embryonic heads from E15.5 mouse embryos were fixed overnight at 4°C with PFA 4%, rinsed with PBS and then cryopreserved through 30% sucrose and frozen in optimal cutting temperature (OCT; Sakura, Torrance, CA, USA). Eyes were cryo-sectioned in 16μm thick sections that were dried 1 hour prior to immunostaining. Immunostaining was done using a Rabbit anti-dsRed (1:1000; Clonetech; # 632496) or a Rabbit anti-Brn3b (Pou4f2) (1:200; Santa Cruz) and images were acquired using an Eclipse 90i epifluorescence microscope (Nikon). All *in situ* hybridizations were retrieved either from the Allen Developing Mouse Brain Atlas (yellow background, www.brain-map.org) or from the digital atlas of gene expression patterns in the mouse (white background images, GenePaint.org).

