## Supplementary material for "Transcriptional logic of cell fate specification and axon guidance in early born retinal neurons revealed by single-cell mRNA profiling"

**KEYWORDS:** Neurogenesis, Retinal development, Single-cell RNA-seq, Retinal ganglion cell, Axon guidance, Cell fate specification

#### SUPPLEMENTARY FIGURE LEGENDS

**Figure S1 — t-SNE reduction of the two replicates from Loupe Cell Browser. A**. t-SNE space of the first replicate (index F2) showing 2675 single transcriptomic profiles, generated with the Loupe Cell Browser software with default parameters. **B**. t-SNE space of the second replicate (index E2) showing 2673 single transcriptomic profiles, generated with the Loupe Cell Browser software with default parameters. Cell types were inferred from marker genes.

**Figure S2 — Gene expression pattern identifying the cell type clustering. A-L**. Feature plots representing the expression patterns of the cycle cycle exit gene (*Top2a*), the neuronal-specific gene (*Btg2*), neuroblast transcription factors (*Neurod4*, *Pax6*), initiation of axonal growth gene (*Pcdh17*), RGC markers (*Pou6f2*, *Elavl4*), amacrine cells (*Prox1*), horizontal cells (*Onecut1*), photoreceptor cells (*Otx2*, *Thrb*) and cone photoreceptors (*Rbp4*) on the t-SNE space, colored by grey to red gradient representing the gene expression levels.

**Figure S3 — Unsupervised clustering validated by ISH. A-F**. ISH of E15.5 eye areas from the Developing Allen Brain Atlas validating the main neuronal clusters with *Otx2* for cone photoreceptors, *Tfap2b* for amacrine cells, *Onecut2* for amacrine/horizontal cells, *Isl1*, *Sncg*, and *Igf1* for RGC cells.

**Figure S4 — The cell cycle organization.** Cells colored by their cell cycle phase score on the UMAP space.

**Figure S5 — Local transcriptomic evolutions by RNA velocity. A.** RNA-velocity plot embedding in the UMAP space the putative near transcription evolutions of each cells on the F2 replicate by the scVelo python package. B. UMAP color-coded by root and end points of the velocity representation on the F2 replicate. C-D. Left parts: Phase portraits of Dlx1 and Prdm1. Right parts: Cell velocity variances for the two genes are represented on an UMAP by color-coding from blue, decreasing to red, increasing the expression.

**Figure S6 — Gene impacting the branching lineage by nodes.** BEAM analysis of each nodes of the pseudotime lineage tree showing the differentially expressed genes between the node emerging cell fates. BEAM heatmap were represented in reduced states to show genes involved with the left fate or the right fate of each nodes. .

**Figure S7 — Expression pattern of miRNAs. A-B**. Feature plots colored by expression levels of miRNAs expressed in the dataset on the t-SNE space. *miR124a-1hg* and *miR124-2hg* shows preferential expression that may trigger AC/HC fates.

**Figure S8 — Morphological identity patterns in function of the t-SNE dimensions for RGC.** Matrix of feature plots of RGC colored by retinal quadrant identity, VT (ventrotemporal), DN (dorsonasal), DT (dorsotemporal), VN (ventronasal), defined by expression of morphological markers genes. Each row and column represent an outputted dimension of the RGC t-SNE space between 1 and 5.

**Figure S9 — Co-expression of ipsilaterally-expressed genes. A-C**. Venn diagram representing the number of RGC cells co-expressing ispsilateral markers in single-cell experiments at E14.5, E15.5 and 16.5 respectively. Majority of RGC cells are not coexpressing ipsilateral markers. **D-F**. Corresponding matrix representing the number of cells expressing the different combinations of the markers. **G**. Cumulative barplots representing the proportion of cells expressing 1, 2, 3 or 4 ipsilateral markers per ages.

**Figure S10 — Expression pattern of major transcription regulators of the Prdm family. A-D**. Feature plots representing the expression patterns of major transcription regulators of the *Prdm* family on the t-SNE space.

**Figure S11 — Expression pattern of metabolic genes *Ldha* and *Ldhb*. A-B**. Feature plots representing the expression patterns of *Ldha* and *Ldhb* on the t-SNE space implying different energetic behavior in function of cell populations.

**Figure S12 — Expression pattern of a potential receptor/ligand/effector couple. A-C**. Feature plots representing the expression patterns of *Adam10*, *Kit* and *Kitl* of the RGC cells on the RGC part of the t-SNE space showing potential interactions of a ligand (*Kitl*) its effector (*Adam10*), and its receptor (*Kit*).

#### SUPPLEMENTARY TABLES LEGENDS

**Table S1 — Differentially expressed genes of the unsupervised clustering reveal cell identities**
Top 15 genes markers for each of the 14 clusters of the merged dataset. The differentially expressed gene analysis was performed with Seurat with default parameters.

**Table S2 — Heterogeneity of the RGC cell waves**
Identity of genes composing each transcriptomic wave of the RGC cells.

**Table S3 — Enrichments in the timed RGC cells clusters**
Markers genes enriched in each of the timed RGC clusters (Young RGC, Mid RGC1, Mid RGC2 and Old RGC)

**Table S4 — Definition of the morphological retinal quadrants**
Morphological genes used to define the ventral, temporal, dorsal and nasal quadrants of the retina.

**Table S5 — Dorsal and ventral genes**
Markers genes enriched in each of the timed RGC clusters.

### Figure S1

A

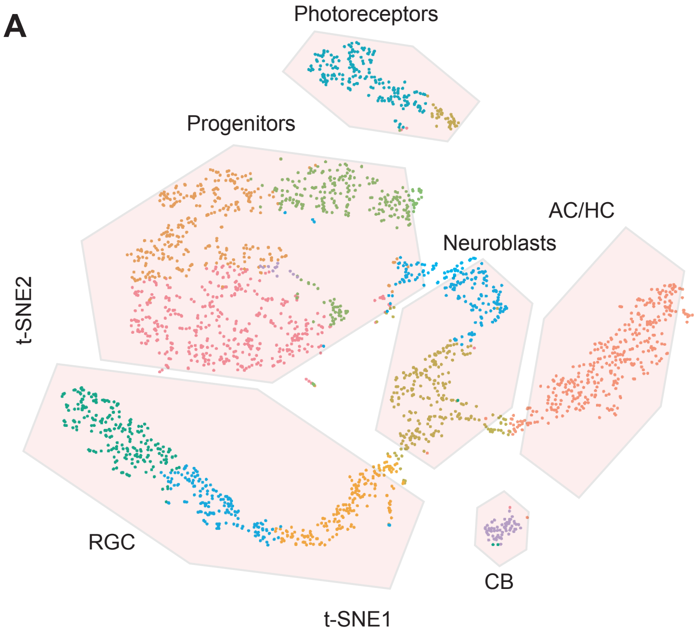
Photoreceptors
Progenitors
AC/HC
Neuroblasts
t-SNE2
RGC
CB
t-SNE1

B

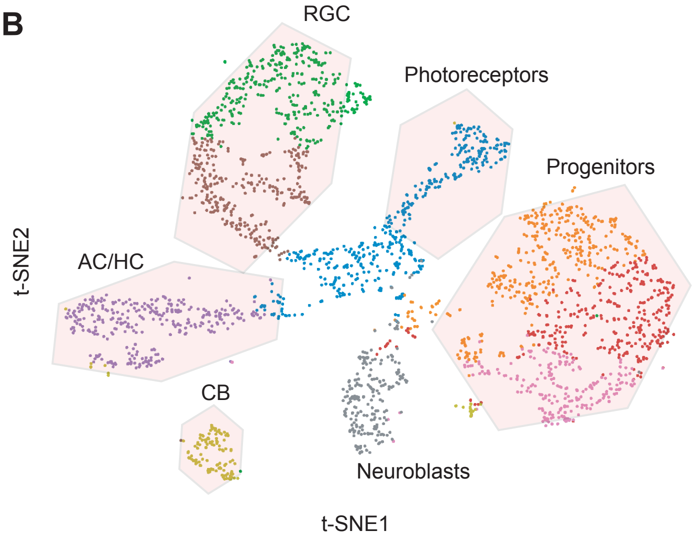
RGC
Photoreceptors
Progenitors
AC/HC
t-SNE2
CB
Neuroblasts
t-SNE1

***Figure S2***

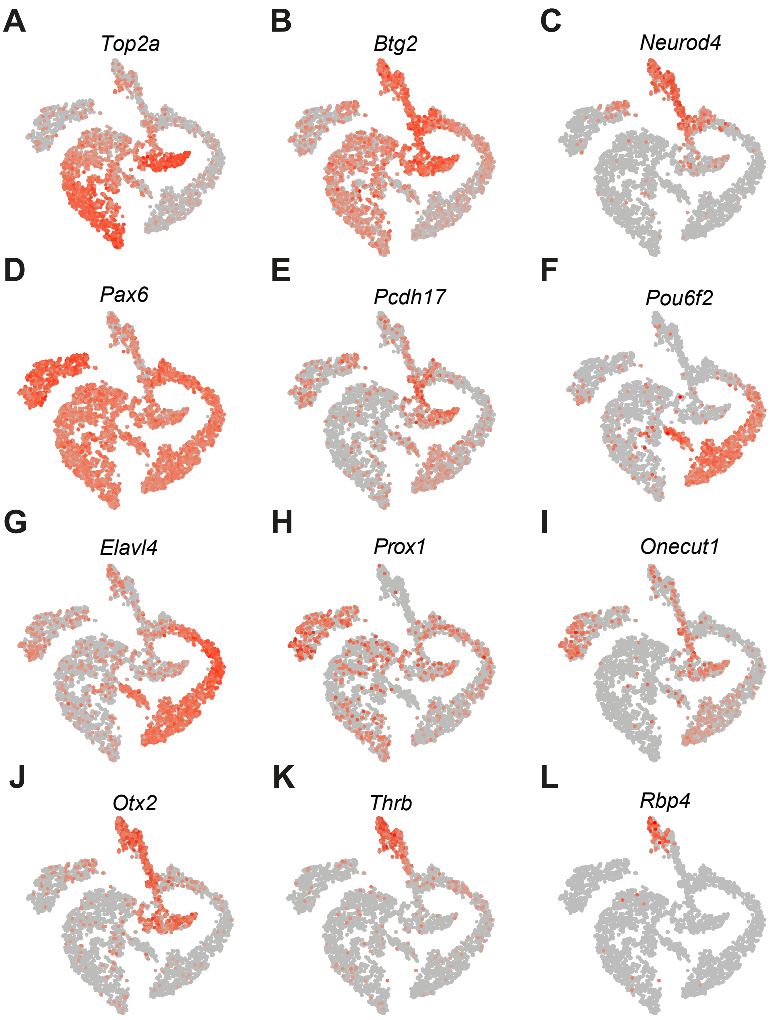
A
Top2a
B
Btg2
C
Neurod4
D
Pax6
E
Pcdh17
F
Pou6f2
G
Elavl4
H
Prox1
I
Onecut1
J
Otx2
K
Thrb
L
Rbp4

*Figure S3*

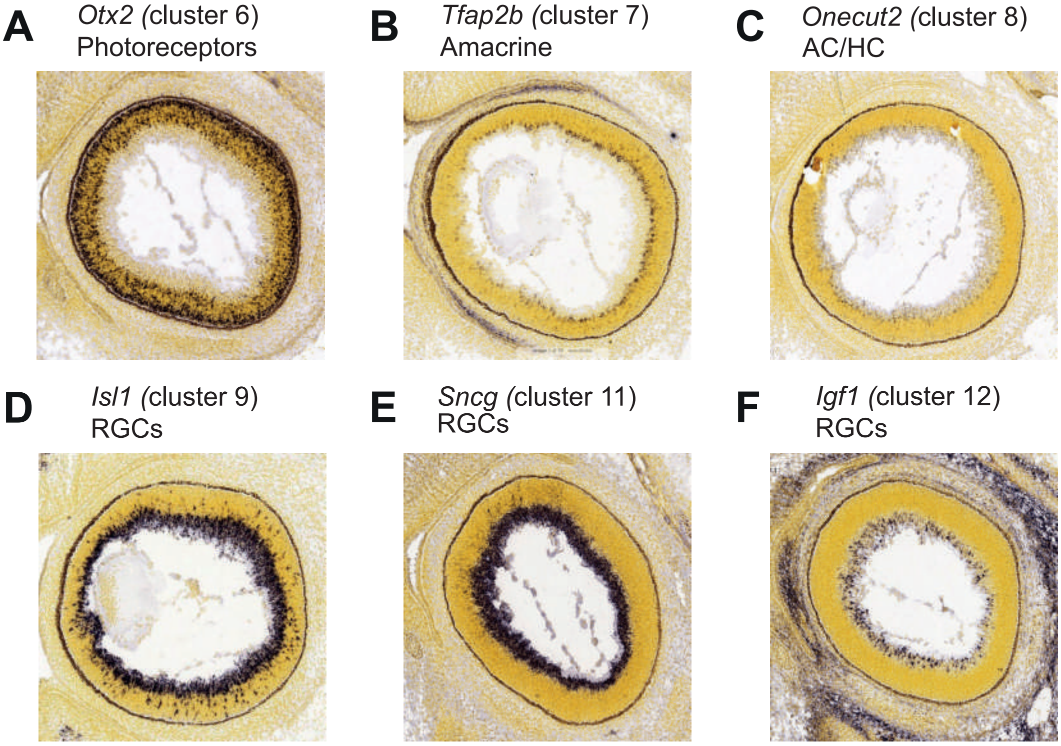
A
*Otx2* (cluster 6)
Photoreceptors
B
*Tfap2b* (cluster 7)
Amacrine
C
*Onecut2* (cluster 8)
AC/HC
D
*Isl1* (cluster 9)
RGCs
E
*Sncg* (cluster 11)
RGCs
F
*Igf1* (cluster 12)
RGCs

*Figure S4*

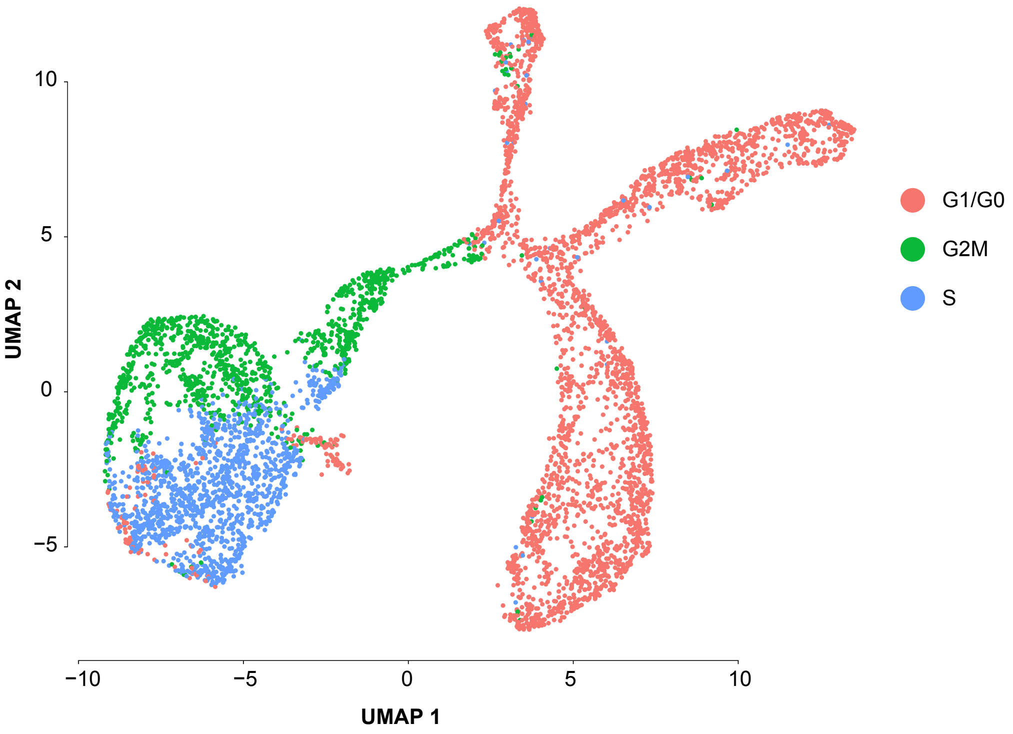

*Figure S5*

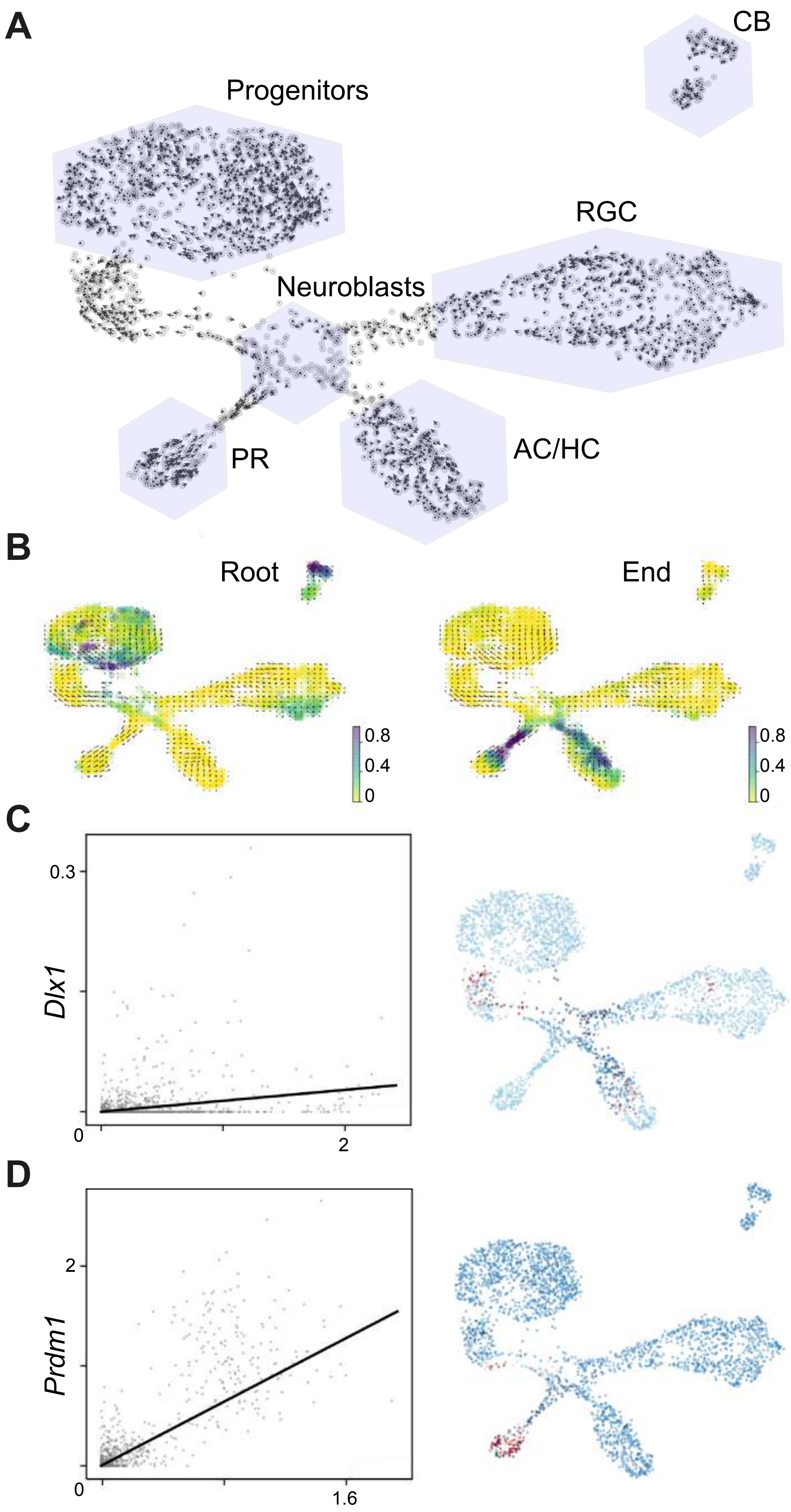

A
CB
Progenitors
RGC
Neuroblasts
PR
AC/HC
B
Root
End
0.8
0.4
0
0.8
0.4
0
C
0.3
Dlx1
0
2
D
2
Prdm1
0
1.6

### Figure S6

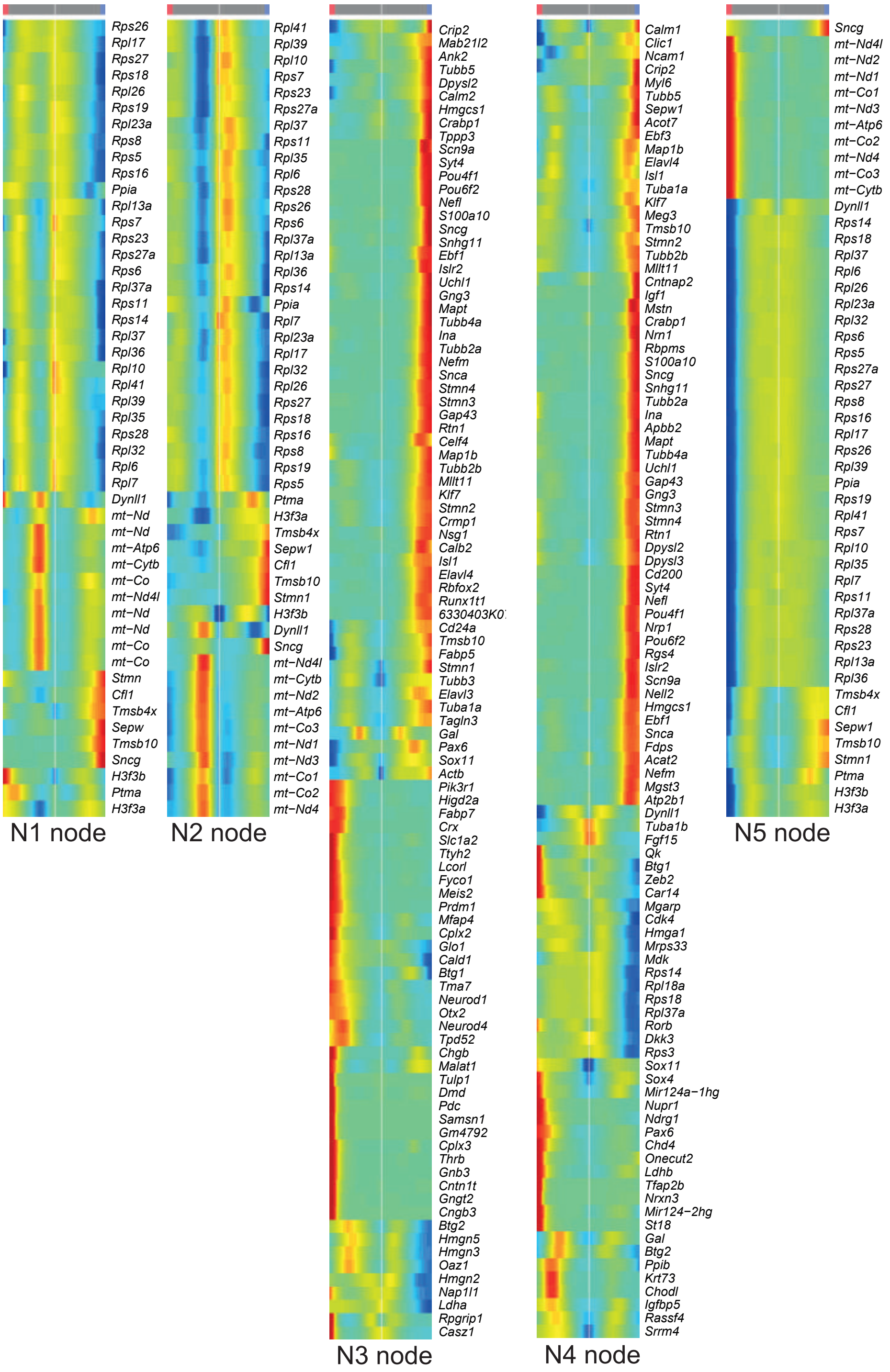
Rps26
Rpl17
Rps27
Rps18
Rpl26
Rps19
Rpl23a
Rps8
Rps5
Rps16
Ppia
Rpl13a
Rps7
Rps23
Rps27a
Rps6
Rpl37a
Rps11
Rps14
Rpl37
Rpl36
Rpl10
Rpl41
Rpl39
Rpl35
Rps28
Rpl32
Rpl6
Rpl7
Dynll1
mt−Nd
mt−Nd
mt−Atp6
mt−Cytb
mt−Co
mt−Nd4l
mt−Nd
mt−Nd
mt−Co
mt−Co
Stmn
Cfl1
Tmsb4x
Sepw
Tmsb10
Sncg
H3f3b
Ptma
H3f3a
N1 node
Rpl41
Rpl39
Rpl10
Rps7
Rps23
Rps27a
Rpl37
Rps11
Rpl35
Rpl6
Rps28
Rps26
Rps6
Rpl37a
Rpl13a
Rpl36
Rps14
Ppia
Rpl7
Rpl23a
Rpl17
Rpl32
Rpl26
Rps27
Rps18
Rps16
Rps8
Rps19
Rps5
Ptma
H3f3a
Tmsb4x
Sepw1
Cfl1
Tmsb10
Stmn1
H3f3b
Dynll1
Sncg
mt−Nd4l
mt−Cytb
mt−Nd2
mt−Atp6
mt−Co3
mt−Nd1
mt−Nd3
mt−Co1
mt−Co2
mt−Nd4
N2 node
Crip2
Mab21l2
Ank2
Tubb5
Dpysl2
Hmgcs1
Crabp1
Tppp3
Scn9a
Syt4
Pou4f1
Pou6f2
Nefl
S100a10
Sncg
Snhg11
Ebf1
Islr2
Uchl1
Gng3
Mapt
Tubb4a
Ina
Tubb2a
Nefm
Snca
Stmn4
Stmn3
Gap43
Rtn1
Celf4
Map1b
Tubb2b
Mllt11
Klf7
Stmn2
Crmp1
Nsg1
Calb2
Isl1
Elavl4
Rbfox2
Runx1t1
6330403K0
Cd24a
Tmsb10
Fabp5
Stmn1
Tubb3
Elavl3
Tuba1a
Tagln3
Gal
Pax6
Sox11
Actb
Pik3r1
Higd2a
Fabp7
Crx
Slc1a2
Ttyh2
Lcorl
Fyco1
Meis2
Prdm1
Mfap4
Cplx2
Glo1
Cald1
Btg1
Tma7
Neurod1
Otx2
Neurod4
Tpd52
Chgb
Malat1
Tulp1
Dmd
Pdc
Samsn1
Gm4792
Cplx3
Thrb
Gnb3
Cntn1t
Gngt2
Cngb3
Btg2
Hmgn5
Hmgn3
Oaz1
Hmgn2
Nap1l1
Ldha
Rpgrip1
Casz1
N3 node
Calm1
Clic1
Ncam1
Crip2
Myl6
Tubb5
Sepw1
Acot7
Ebf3
Map1b
Elavl4
Isl1
Tuba1a
Klf7
Meg3
Tmsb10
Stmn2
Tubb2b
Mllt11
Cntnap2
Igf1
Mstn
Crabp1
Nrn1
Rbpms
S100a10
Sncg
Snhg11
Tubb2a
Ina
Apbb2
Mapt
Tubb4a
Uchl1
Gap43
Gng3
Stmn3
Stmn4
Rtn1
Dpysl2
Dpysl3
Cd200
Syt4
Nefl
Pou4f1
Nrp1
Pou6f2
Rgs4
Islr2
Scn9a
Nell2
Hmgcs1
Ebf1
Snca
Fdps
Acat2
Nefm
Mgst3
Atp2b1
Dynll1
Tuba1b
Fgf15
Qk
Btg1
Zeb2
Car14
Mgarp
Cdk4
Hmga1
Mrps33
Mdk
Rps14
Rpl18a
Rps18
Rpl37a
Rorb
Dkk3
Rps3
Sox11
Sox4
Mir124a−1hg
Nupr1
Ndrg1
Pax6
Chd4
Onecut2
Ldhb
Tfap2b
Nrxn3
Mir124−2hg
St18
Gal
Btg2
Ppib
Krt73
Chodl
Igfbp5
Rassf4
Srrm4
N4 node
Sncg
mt−Nd4l
mt−Nd2
mt−Nd1
mt−Co1
mt−Nd3
mt−Atp6
mt−Co2
mt−Nd4
mt−Co3
mt−Cytb
Dynll1
Rps14
Rps18
Rpl37
Rpl6
Rpl26
Rpl23a
Rpl32
Rps6
Rps5
Rps27a
Rps27
Rps8
Rps16
Rpl17
Rps26
Rpl39
Ppia
Rps19
Rpl41
Rps7
Rpl10
Rpl35
Rpl7
Rps11
Rpl37a
Rps28
Rps23
Rpl13a
Rpl36
Tmsb4x
Cfl1
Sepw1
Tmsb10
Stmn1
Ptma
H3f3b
H3f3a
N5 node

*Figure S7*

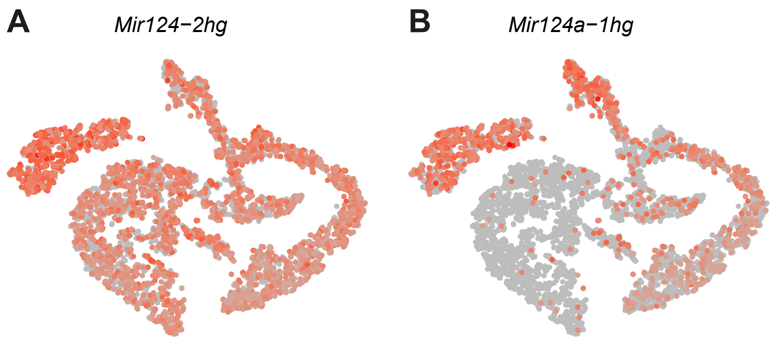

*Figure S8*

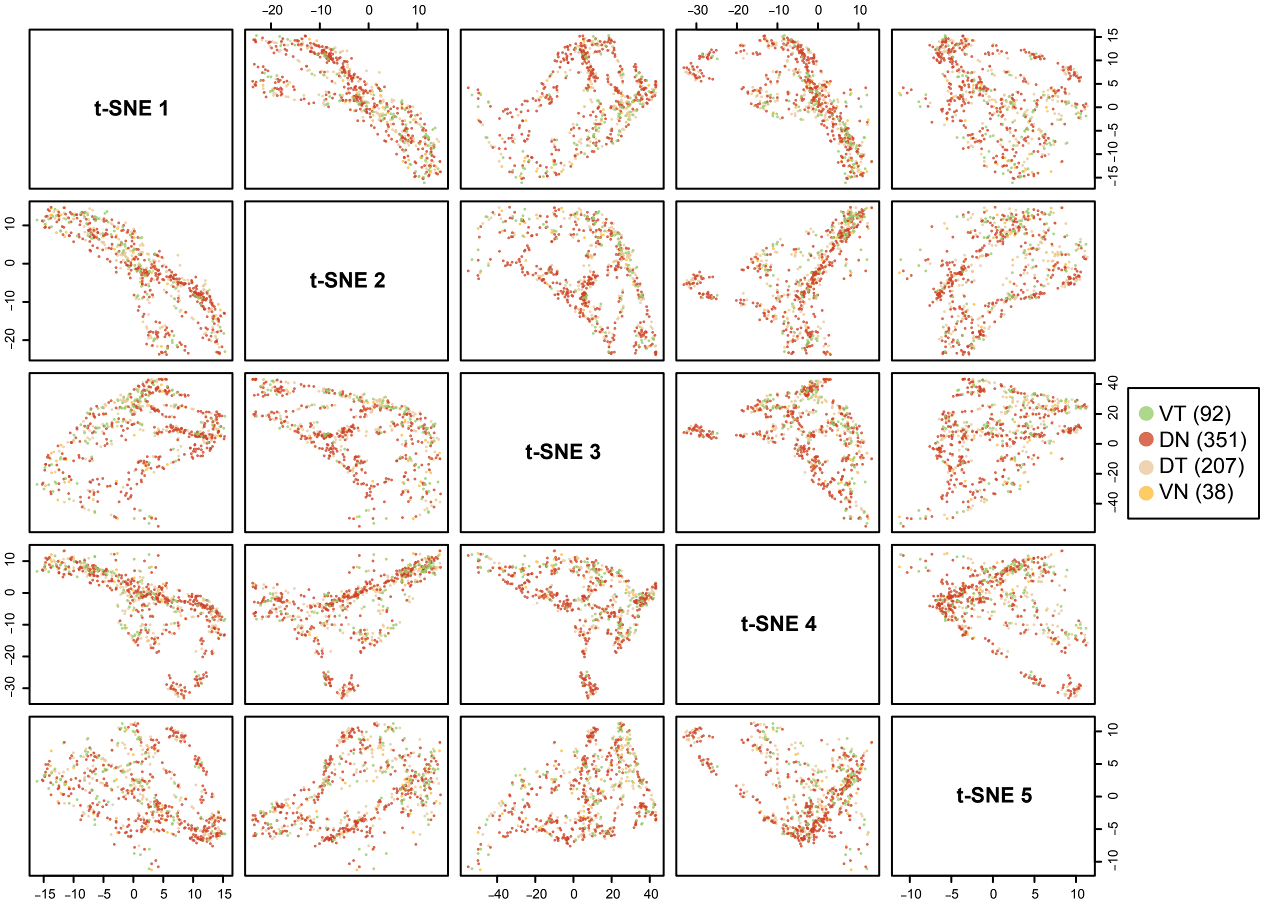

### Figure S9

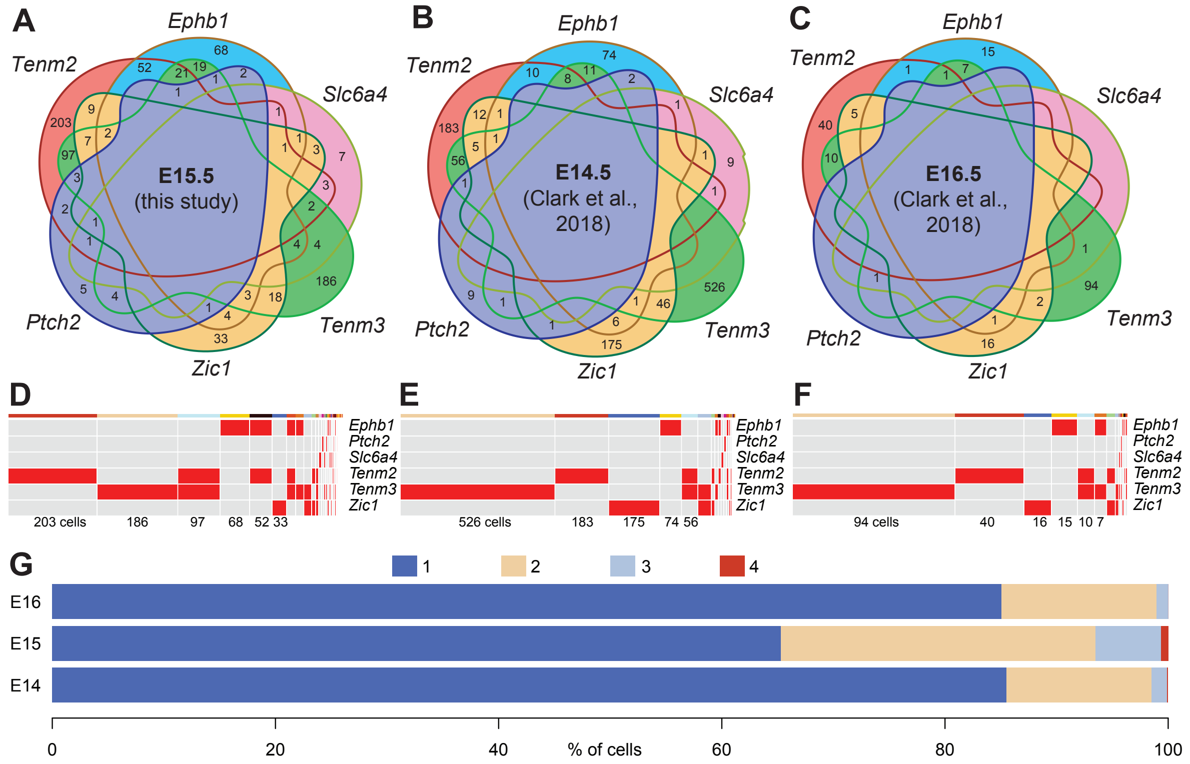
A
B
C
Ephb1
Tenm2
Slc6a4
Ptch2
Tenm3
Zic1
E15.5
(this study)
E14.5
(Clark et al., 2018)
E16.5
(Clark et al., 2018)
D
E
F
203 cells
186
97
68
52 33
526 cells
183
175
74 56
94 cells
40
16
15
10 7
G
1
2
3
4
E16
E15
E14
0
20
40
60
80
100
% of cells

*Figure S10*

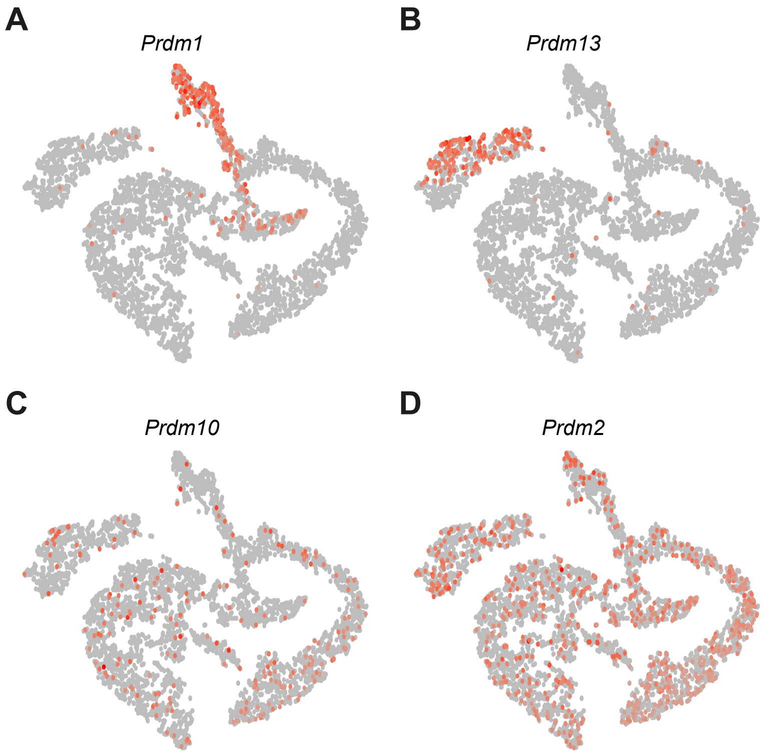

*Figure S11*

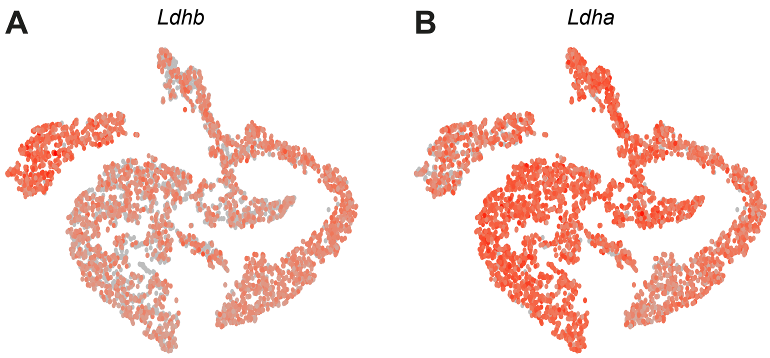

*Figure S12*

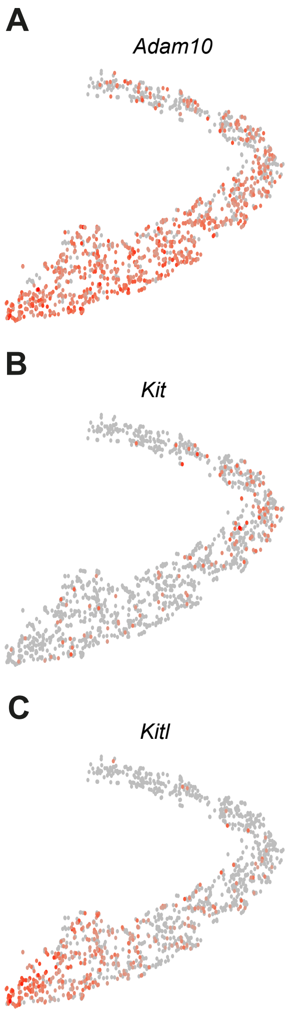

### Table S1

Cluster 0

| gene | p_val | avg_logFC | p_val_adj | cluster |
|---|---|---|---|---|
| *Ccnd1* | 3.57E-173 | 1.0236045 | 5.42E-169 | 0 |
| *Dapl1* | 8.36E-166 | 0.9682639 | 1.27E-161 | 0 |
| *Itm2a* | 5.43E-149 | 0.9596223 | 8.24E-145 | 0 |
| *Cdca7* | 3.11E-177 | 0.9401031 | 4.72E-173 | 0 |
| *Fos* | 4.18E-96 | 0.9302969 | 6.35E-92 | 0 |
| *Mcm3* | 1.08E-159 | 0.9128558 | 1.64E-155 | 0 |
| *Mdk* | 7.15E-164 | 0.8934379 | 1.09E-159 | 0 |
| *Zfp36l1* | 3.36E-157 | 0.8512779 | 5.10E-153 | 0 |
| *Id3* | 9.69E-93 | 0.8099441 | 1.47E-88 | 0 |
| *Sparc* | 3.74E-117 | 0.8044286 | 5.68E-113 | 0 |
| *Mcm4* | 8.17E-122 | 0.795783 | 1.24E-117 | 0 |
| *Mcm5* | 1.78E-113 | 0.7628838 | 2.70E-109 | 0 |
| *Slc2a1* | 9.83E-105 | 0.7584338 | 1.49E-100 | 0 |
| *Serpinh1* | 1.16E-109 | 0.756397 | 1.75E-105 | 0 |
| *Crabp2* | 1.41E-89 | 0.7435662 | 2.15E-85 | 0 |

Cluster 1

| gene | p_val | avg_logFC | p_val_adj | cluster |
|---|---|---|---|---|
| *Hes1* | 1.34E-69 | 1.0849164 | 2.03E-65 | 1 |
| *Ccnd1* | 4.54E-78 | 0.9841358 | 6.90E-74 | 1 |
| *Mcm3* | 1.44E-65 | 0.9431507 | 2.18E-61 | 1 |
| *Fgf15* | 4.67E-71 | 0.933091 | 7.09E-67 | 1 |
| *Ndufa4l2* | 1.40E-34 | 0.9315887 | 2.12E-30 | 1 |
| *Lig1* | 3.28E-51 | 0.8965418 | 4.97E-47 | 1 |
| *Cp* | 1.94E-51 | 0.8936894 | 2.94E-47 | 1 |
| *Slc2a1* | 5.28E-53 | 0.8883581 | 8.01E-49 | 1 |
| *Itm2a* | 6.27E-51 | 0.8802141 | 9.51E-47 | 1 |
| *Dapl1* | 1.93E-64 | 0.8802139 | 2.93E-60 | 1 |
| *Slc1a3* | 5.05E-52 | 0.8710798 | 7.66E-48 | 1 |
| *Mdk* | 2.10E-65 | 0.8648439 | 3.19E-61 | 1 |
| *Serpinh1* | 6.72E-47 | 0.8522818 | 1.02E-42 | 1 |
| *Col9a1* | 9.86E-37 | 0.8363604 | 1.50E-32 | 1 |
| *Id3* | 1.75E-47 | 0.8044184 | 2.66E-43 | 1 |

Cluster 2

| gene | p_val | avg_logFC | p_val_adj | cluster |
|---|---|---|---|---|
| *Fgf15* | 1.34E-260 | 1.5417718 | 2.04E-256 | 2 |
| *Hist1h1b* | 0 | 1.518998 | 0 | 2 |
| *2810417H1* | 8.48E-290 | 1.4801442 | 1.29E-285 | 2 |
| *Fos* | 2.35E-149 | 1.2748731 | 3.56E-145 | 2 |
| *Top2a* | 4.26E-255 | 1.1705807 | 6.47E-251 | 2 |
| *Hes1* | 1.16E-197 | 1.1613276 | 1.76E-193 | 2 |
| *Id2* | 1.81E-172 | 1.0830062 | 2.75E-168 | 2 |
| *Id3* | 3.51E-162 | 1.0553903 | 5.32E-158 | 2 |
| *Hist1h4d* | 8.79E-165 | 1.0407038 | 1.33E-160 | 2 |
| *Zfp36l1* | 2.58E-188 | 1.0272495 | 3.91E-184 | 2 |
| *Esco2* | 2.65E-281 | 1.0115413 | 4.02E-277 | 2 |
| *Ccnd1* | 1.20E-165 | 0.9883776 | 1.82E-161 | 2 |
| *Spc25* | 2.68E-221 | 0.985421 | 4.07E-217 | 2 |
| *Rrm2* | 2.55E-140 | 0.94441 | 3.87E-136 | 2 |
| *Lig1* | 7.20E-143 | 0.9273175 | 1.09E-138 | 2 |

Cluster 3

| gene | p_val | avg_logFC | p_val_adj | cluster |
|---|---|---|---|---|
| *Cenpf* | 1.93E-256 | 1.6697339 | 2.94E-252 | 3 |
| *Ccnb1* | 1.03E-292 | 1.6314366 | 1.56E-288 | 3 |
| *Ube2c* | 5.72E-246 | 1.5592696 | 8.68E-242 | 3 |
| *Cks2* | 5.56E-212 | 1.5154302 | 8.44E-208 | 3 |
| *Cenpa* | 8.72E-246 | 1.4453531 | 1.32E-241 | 3 |
| *Cdc20* | 4.14E-190 | 1.3837907 | 6.28E-186 | 3 |
| *Prc1* | 2.27E-139 | 1.3787874 | 3.44E-135 | 3 |
| *Birc5* | 2.28E-234 | 1.3572867 | 3.47E-230 | 3 |
| *Cenpe* | 7.04E-238 | 1.2929958 | 1.07E-233 | 3 |
| *Nusap1* | 7.21E-163 | 1.2566128 | 1.09E-158 | 3 |
| *Tpx2* | 1.61E-228 | 1.2522382 | 2.45E-224 | 3 |
| *Cdca8* | 6.55E-198 | 1.2439556 | 9.94E-194 | 3 |
| *Arl6ip1* | 1.29E-102 | 1.2100156 | 1.96E-98 | 3 |
| *Hmmr* | 1.05E-221 | 1.1998652 | 1.60E-217 | 3 |
| *Knstrn* | 8.46E-239 | 1.1969341 | 1.28E-234 | 3 |

Cluster 4

| gene | p_val | avg_logFC | p_val_adj | cluster |
|---|---|---|---|---|
| *Psip1* | 1.85E-155 | 1.720335 | 2.80E-151 | 4 |
| *Top2a* | 9.12E-224 | 1.704728 | 1.38E-219 | 4 |
| *Penk* | 0 | 1.5352682 | 0 | 4 |
| *Pbk* | 1.47E-246 | 1.4654442 | 2.24E-242 | 4 |
| *Cenpf* | 1.72E-183 | 1.4559108 | 2.61E-179 | 4 |
| *Nusap1* | 3.00E-228 | 1.434909 | 4.55E-224 | 4 |
| *Rrm2* | 4.79E-168 | 1.4102074 | 7.27E-164 | 4 |
| *Cks2* | 5.87E-171 | 1.3722483 | 8.91E-167 | 4 |
| *Hist1h1b* | 3.04E-163 | 1.3698736 | 4.61E-159 | 4 |
| *Hmgb2* | 1.39E-177 | 1.3535926 | 2.11E-173 | 4 |
| *Ccnb1* | 2.47E-175 | 1.340922 | 3.75E-171 | 4 |
| *Cdk1* | 5.41E-194 | 1.3404176 | 8.21E-190 | 4 |
| *Ube2c* | 1.32E-136 | 1.3317928 | 2.00E-132 | 4 |
| *Tpx2* | 3.89E-181 | 1.3282019 | 5.90E-177 | 4 |
| *Cenpe* | 2.37E-171 | 1.3221295 | 3.60E-167 | 4 |

Cluster 5

| gene | p_val | avg_logFC | p_val_adj | cluster |
|---|---|---|---|---|
| *Gal* | 2.53E-10 | 1.3649989 | 3.83E-06 | 5 |
| *Pcdh17* | 3.37E-71 | 1.3585195 | 5.12E-67 | 5 |
| *Btg2* | 7.71E-137 | 1.3540663 | 1.17E-132 | 5 |
| *Psip1* | 1.29E-67 | 1.134719 | 1.95E-63 | 5 |
| *Gadd45a* | 2.02E-123 | 1.0962028 | 3.06E-119 | 5 |
| *Mgarp* | 1.34E-56 | 1.0645597 | 2.03E-52 | 5 |
| *Rassf4* | 3.80E-104 | 0.9992351 | 5.77E-100 | 5 |
| *Mfng* | 4.76E-94 | 0.922558 | 7.22E-90 | 5 |
| *Hmgn3* | 2.18E-78 | 0.9195723 | 3.30E-74 | 5 |
| *Hmgn5* | 6.04E-64 | 0.8589627 | 9.17E-60 | 5 |
| *Rgs16* | 1.48E-84 | 0.8272245 | 2.25E-80 | 5 |
| *Srrm4* | 1.40E-61 | 0.8180129 | 2.13E-57 | 5 |
| *Neurod4* | 8.18E-64 | 0.7990576 | 1.24E-59 | 5 |
| *Pkib* | 2.86E-89 | 0.7961818 | 4.33E-85 | 5 |
| *Notch1* | 3.62E-45 | 0.7400647 | 5.50E-41 | 5 |

Cluster 6

| gene | p_val | avg_logFC | p_val_adj | cluster |
|---|---|---|---|---|
| *Gngt2* | 0 | 2.8853448 | 0 | 6 |
| *Fabp7* | 8.50E-245 | 2.5623106 | 1.29E-240 | 6 |
| *Thrb* | 0 | 2.497383 | 0 | 6 |
| *Neurod1* | 0 | 2.3671284 | 0 | 6 |
| *Meis2* | 0 | 2.2014893 | 0 | 6 |
| *Pdc* | 0 | 2.2009538 | 0 | 6 |
| *Pik3r1* | 1.15E-284 | 2.2007679 | 1.75E-280 | 6 |
| *Neurod4* | 0 | 2.1128351 | 0 | 6 |
| *Otx2* | 0 | 1.889568 | 0 | 6 |
| *Gnb3* | 0 | 1.7904282 | 0 | 6 |
| *Crx* | 0 | 1.7630776 | 0 | 6 |
| *Mfap4* | 8.24E-232 | 1.6888914 | 1.25E-227 | 6 |
| *Chgb* | 3.54E-161 | 1.5858375 | 5.38E-157 | 6 |
| *Cplx2* | 4.39E-175 | 1.5774599 | 6.66E-171 | 6 |
| *Samsn1* | 0 | 1.5323024 | 0 | 6 |

Cluster 7

| gene | p_val | avg_logFC | p_val_adj | cluster |
|---|---|---|---|---|
| *Nrxn3* | 9.20E-307 | 1.4577913 | 1.40E-302 | 7 |
| *Tfap2b* | 0 | 1.4494792 | 0 | 7 |
| *Igfbp5* | 1.41E-251 | 1.4172303 | 2.13E-247 | 7 |
| *Nupr1* | 0 | 1.4168169 | 0 | 7 |
| *Krt73* | 1.78E-241 | 1.3910059 | 2.70E-237 | 7 |
| *Sox11* | 6.74E-218 | 1.3321591 | 1.02E-213 | 7 |
| *St18* | 0 | 1.3218674 | 0 | 7 |
| *Chodl* | 4.48E-281 | 1.2899239 | 6.79E-277 | 7 |
| *Cdkn1c* | 1.43E-84 | 1.2883829 | 2.17E-80 | 7 |
| *Sox4* | 2.26E-178 | 1.23033 | 3.43E-174 | 7 |
| *Ndrg1* | 6.66E-200 | 1.1820073 | 1.01E-195 | 7 |
| *Carhsp1* | 9.60E-161 | 1.1593832 | 1.46E-156 | 7 |
| *Calb2* | 3.24E-90 | 1.1299581 | 4.92E-86 | 7 |
| *H19* | 1.44E-22 | 1.0936732 | 2.19E-18 | 7 |
| *Lbh* | 3.77E-121 | 1.0682108 | 5.71E-117 | 7 |

Cluster 8

| gene | p_val | avg_logFC | p_val_adj | cluster |
|---|---|---|---|---|
| *Tfap2b* | 0 | 2.2313767 | 0 | 8 |
| *Onecut2* | 5.31E-137 | 1.9097552 | 8.05E-133 | 8 |
| *Nnat* | 5.65E-67 | 1.8528057 | 8.58E-63 | 8 |
| *Nrxn3* | 3.25E-244 | 1.6890694 | 4.93E-240 | 8 |
| *Celf4* | 5.49E-130 | 1.4480621 | 8.34E-126 | 8 |
| *Pmfbp1* | 0 | 1.269631 | 0 | 8 |
| *A930011G2* | 6.30E-128 | 1.1924201 | 9.56E-124 | 8 |
| *Pcbp3* | 4.24E-90 | 1.1801708 | 6.44E-86 | 8 |
| *Ldhb* | 1.08E-83 | 1.1551818 | 1.64E-79 | 8 |
| *Sox4* | 2.37E-84 | 1.1081805 | 3.60E-80 | 8 |
| *Rnd3* | 1.07E-53 | 1.0107146 | 1.63E-49 | 8 |
| *Auts2* | 1.10E-59 | 0.9449796 | 1.66E-55 | 8 |
| *Runx1t1* | 1.90E-38 | 0.9136571 | 2.88E-34 | 8 |
| *Dcx* | 7.64E-42 | 0.9080534 | 1.16E-37 | 8 |
| *Bcl11a* | 1.72E-48 | 0.8930283 | 2.61E-44 | 8 |

Cluster 9

| gene | p_val | avg_logFC | p_val_adj | cluster |
|---|---|---|---|---|
| *Isl1* | 3.91E-193 | 1.5788354 | 5.94E-189 | 9 |
| *Elavl4* | 2.74E-169 | 1.482858 | 4.15E-165 | 9 |
| *Ccer2* | 3.99E-164 | 1.1697411 | 6.06E-160 | 9 |
| *Pou4f2* | 1.68E-212 | 1.1414261 | 2.56E-208 | 9 |
| *Tubb3* | 2.77E-126 | 1.0910579 | 4.20E-122 | 9 |
| *Gal* | 1.20E-13 | 1.0045128 | 1.83E-09 | 9 |
| *Rassf4* | 2.25E-106 | 0.9431221 | 3.42E-102 | 9 |
| *Chrna3* | 6.47E-96 | 0.9170162 | 9.82E-92 | 9 |
| *Klf7* | 9.70E-89 | 0.891623 | 1.47E-84 | 9 |
| *Mfng* | 2.28E-126 | 0.8894969 | 3.46E-122 | 9 |
| *Map1b* | 4.02E-82 | 0.877401 | 6.10E-78 | 9 |
| *Snca* | 2.09E-66 | 0.8462842 | 3.18E-62 | 9 |
| *Miat* | 1.29E-88 | 0.844016 | 1.95E-84 | 9 |
| *Elavl3* | 1.67E-85 | 0.823252 | 2.53E-81 | 9 |
| *Cntn2* | 2.28E-91 | 0.811699 | 3.46E-87 | 9 |

Cluster 10

| gene | p_val | avg_logFC | p_val_adj | cluster |
|---|---|---|---|---|
| *Gap43* | 1.11E-193 | 1.76921 | 1.68E-189 | 10 |
| *Sncg* | 6.71E-183 | 1.607325 | 1.02E-178 | 10 |
| *Nefm* | 7.24E-229 | 1.6039133 | 1.10E-224 | 10 |
| *Nefl* | 2.51E-188 | 1.5879133 | 3.81E-184 | 10 |
| *S100a10* | 2.96E-211 | 1.4971431 | 4.49E-207 | 10 |
| *Snca* | 4.37E-203 | 1.3960487 | 6.63E-199 | 10 |
| *Isl1* | 1.53E-198 | 1.3588494 | 2.32E-194 | 10 |
| *Elavl4* | 3.01E-202 | 1.3551689 | 4.57E-198 | 10 |
| *Stmn2* | 5.15E-169 | 1.3287006 | 7.82E-165 | 10 |
| *Gng3* | 1.95E-208 | 1.3160503 | 2.97E-204 | 10 |
| *Akap7* | 1.02E-270 | 1.2520231 | 1.55E-266 | 10 |
| *Crabp1* | 1.71E-15 | 1.216465 | 2.59E-11 | 10 |
| *Mllt11* | 4.80E-162 | 1.2058069 | 7.28E-158 | 10 |
| *Apbb2* | 1.00E-169 | 1.2037585 | 1.52E-165 | 10 |
| *Klf7* | 1.70E-173 | 1.1906561 | 2.58E-169 | 10 |

Cluster 11

| gene | p_val | avg_logFC | p_val_adj | cluster |
|---|---|---|---|---|
| *Dpysl2* | 3.75E-201 | 1.673083 | 5.69E-197 | 11 |
| *Sncg* | 2.47E-189 | 1.6644576 | 3.75E-185 | 11 |
| *Pou4f1* | 1.24E-233 | 1.636452 | 1.88E-229 | 11 |
| *Gap43* | 1.86E-167 | 1.5953867 | 2.82E-163 | 11 |
| *Nefl* | 3.34E-158 | 1.4271139 | 5.07E-154 | 11 |
| *Ina* | 9.06E-212 | 1.383574 | 1.37E-207 | 11 |
| *S100a10* | 7.47E-187 | 1.359117 | 1.13E-182 | 11 |
| *Gnb1* | 1.94E-177 | 1.3452769 | 2.94E-173 | 11 |
| *Nefm* | 6.33E-180 | 1.3172435 | 9.60E-176 | 11 |
| *Mapt* | 4.52E-200 | 1.2832422 | 6.86E-196 | 11 |
| *Actb* | 5.80E-176 | 1.2607375 | 8.80E-172 | 11 |
| *Runx1t1* | 3.36E-186 | 1.25489 | 5.10E-182 | 11 |
| *Snhg11* | 3.83E-152 | 1.2480071 | 5.82E-148 | 11 |
| *Uchl1* | 2.47E-166 | 1.2354923 | 3.75E-162 | 11 |
| *Pou6f2* | 3.66E-216 | 1.2332414 | 5.55E-212 | 11 |

Cluster 12

| gene | p_val | avg_logFC | p_val_adj | cluster |
|---|---|---|---|---|
| *Sncg* | 5.06E-200 | 1.8778599 | 7.68E-196 | 12 |
| *Snhg11* | 3.97E-269 | 1.8495724 | 6.02E-265 | 12 |
| *Tubb2a* | 1.69E-205 | 1.6705797 | 2.56E-201 | 12 |
| *Mapt* | 6.91E-237 | 1.5932211 | 1.05E-232 | 12 |
| *Meg3* | 6.60E-161 | 1.5553131 | 1.00E-156 | 12 |
| *Nefl* | 2.75E-168 | 1.5041275 | 4.18E-164 | 12 |
| *Gap43* | 4.91E-137 | 1.4559236 | 7.45E-133 | 12 |
| *Igf1* | 2.68E-298 | 1.4351284 | 4.07E-294 | 12 |
| *Syt4* | 3.12E-258 | 1.4351217 | 4.74E-254 | 12 |
| *Stmn3* | 1.56E-181 | 1.4299666 | 2.36E-177 | 12 |
| *Ina* | 6.05E-209 | 1.4239091 | 9.18E-205 | 12 |
| *Tubb2b* | 1.18E-162 | 1.4133299 | 1.79E-158 | 12 |
| *Pou6f2* | 1.91E-239 | 1.4033886 | 2.90E-235 | 12 |
| *Uchl1* | 1.75E-179 | 1.3778301 | 2.66E-175 | 12 |
| *S100a10* | 6.79E-170 | 1.2996108 | 1.03E-165 | 12 |

Cluster 13

| gene | p_val | avg_logFC | p_val_adj | cluster |
|---|---|---|---|---|
| *Ebf1* | 2.16E-100 | 1.8645756 | 3.28E-96 | 13 |
| *mt-Nd1* | 1.46E-67 | 1.8153384 | 2.21E-63 | 13 |
| *mt-Nd4* | 5.60E-70 | 1.7221216 | 8.51E-66 | 13 |
| *mt-Co3* | 6.45E-69 | 1.687274 | 9.79E-65 | 13 |
| *mt-Cytb* | 2.11E-67 | 1.6523162 | 3.20E-63 | 13 |
| *mt-Co2* | 3.22E-67 | 1.5992198 | 4.88E-63 | 13 |
| *mt-Atp6* | 5.06E-65 | 1.5814611 | 7.68E-61 | 13 |
| *Stmn4* | 9.42E-42 | 1.5773032 | 1.43E-37 | 13 |
| *Islr2* | 9.51E-64 | 1.5710112 | 1.44E-59 | 13 |
| *Syt4* | 1.39E-52 | 1.5317412 | 2.10E-48 | 13 |
| *mt-Co1* | 2.31E-69 | 1.5118989 | 3.51E-65 | 13 |
| *Stmn3* | 2.82E-41 | 1.4418527 | 4.28E-37 | 13 |
| *Scn9a* | 5.87E-68 | 1.431588 | 8.91E-64 | 13 |
| *mt-Nd3* | 7.91E-41 | 1.4144281 | 1.20E-36 | 13 |
| *Tubb2a* | 1.62E-55 | 1.4081109 | 2.47E-51 | 13 |

### *Table S2*

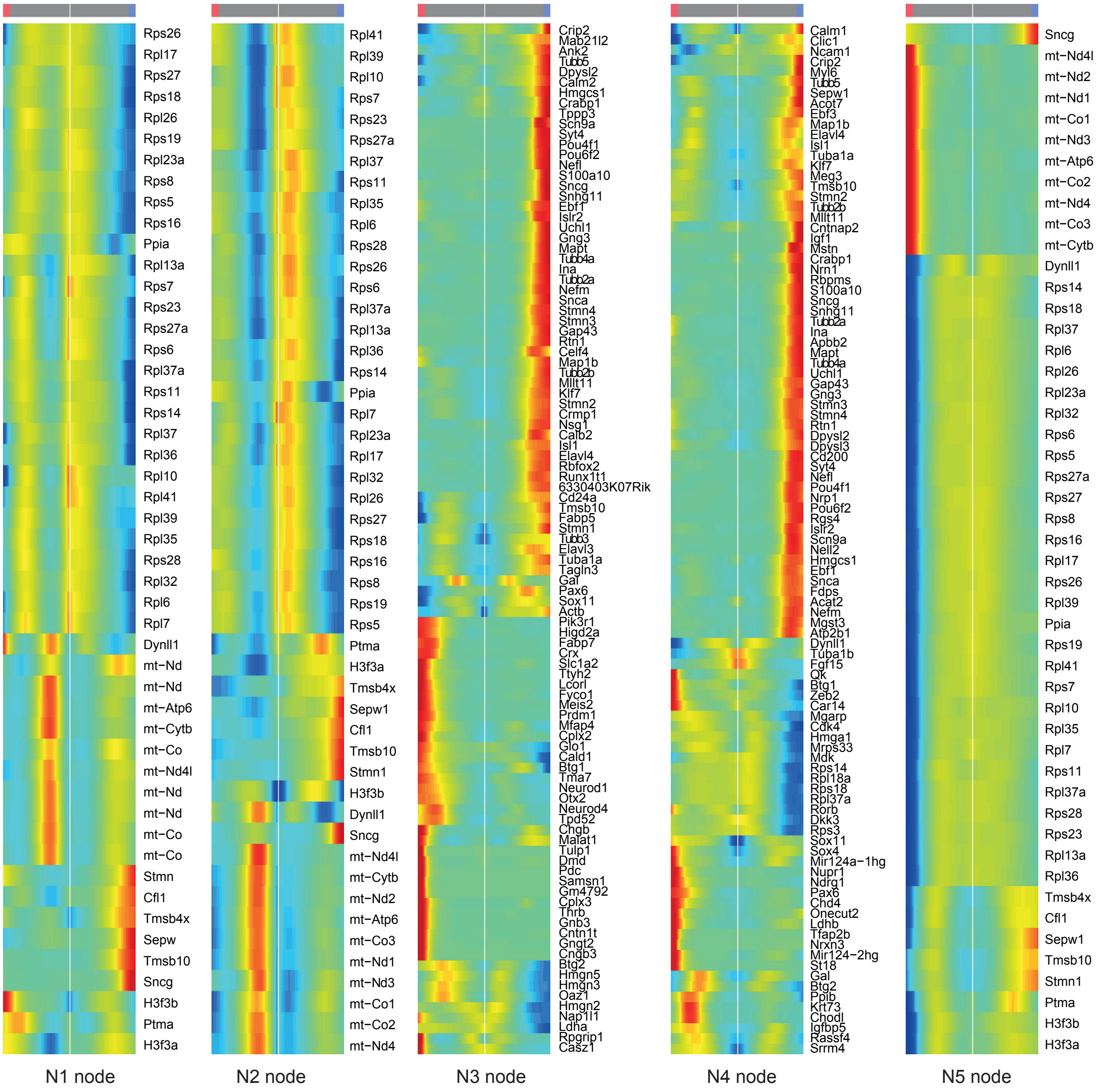
Rps26
Rpl17
Rps27
Rps18
Rpl26
Rps19
Rpl23a
Rps8
Rps5
Rps16
Ppia
Rpl13a
Rps7
Rps23
Rps27a
Rps6
Rpl37a
Rps11
Rps14
Rpl37
Rpl36
Rpl10
Rpl41
Rpl39
Rpl35
Rps28
Rpl32
Rpl6
Rpl7
Dynll1
mt−Nd
mt−Nd
mt−Atp6
mt−Cytb
mt−Co
mt−Nd4l
mt−Nd
mt−Nd
mt−Co
mt−Co
Stmn
Cfl1
Tmsb4x
Sepw
Tmsb10
Sncg
H3f3b
Ptma
H3f3a
N1 node
Rpl41
Rpl39
Rpl10
Rps7
Rps23
Rps27a
Rpl37
Rps11
Rpl35
Rpl6
Rps28
Rps26
Rps6
Rpl37a
Rpl13a
Rpl36
Rps14
Ppia
Rpl7
Rpl23a
Rpl17
Rpl32
Rpl26
Rps27
Rps18
Rps16
Rps8
Rps19
Rps5
Ptma
H3f3a
Tmsb4x
Sepw1
Cfl1
Tmsb10
Stmn1
H3f3b
Dynll1
Sncg
mt−Nd4l
mt−Cytb
mt−Nd2
mt−Atp6
mt−Co3
mt−Nd1
mt−Nd3
mt−Co1
mt−Co2
mt−Nd4
N2 node
Crip2
Mab21l2
Ank2
Tubb5
Dpysl2
Calm2
Hmgcs1
Crabp1
Tppp3
Scn9a
Syt4
Pou4f1
Pou6f2
Nefl
S100a10
Sncg
Snhg11
Ebf1
Islr2
Uchl1
Gng3
Mapt
Tubb4a
Ina
Tubb2a
Nefm
Snca
Stmn4
Stmn3
Gap43
Rtn1
Celf4
Map1b
Tubb2b
Mllt11
Klf7
Stmn2
Crmp1
Nsg1
Calb2
Isl1
Elavl4
Rbfox2
Runx1t1
6330403K07Rik
Cd24a
Tmsb10
Fabp5
Stmn1
Tubb3
Elavl3
Tuba1a
Tagln3
Gal
Pax6
Sox11
Actb
Pik3r1
Higd2a
Fabp7
Crx
Slc1a2
Ttyh2
Lcorl
Fyco1
Meis2
Prdm1
Mfap4
Cplx2
Glo1
Cald1
Btg1
Tma7
Neurod1
Otx2
Neurod4
Tpd52
Chgb
Malat1
Tulp1
Dmd
Pdc
Samsn1
Gm4792
Cplx3
Thrb
Gnb3
Cntn1t
Gngt2
Cngb3
Btg2
Hmgn5
Hmgn3
Oaz1
Hmgn2
Nap1l1
Ldha
Rpgrip1
Casz1
N3 node
Calm1
Clic1
Ncam1
Crip2
Myl6
Tubb5
Sepw1
Acot7
Ebf3
Map1b
Elavl4
Isl1
Tuba1a
Klf7
Meg3
Tmsb10
Stmn2
Tubb2b
Mllt11
Cntnap2
Igf1
Mstn
Crabp1
Nrn1
Rbpms
S100a10
Sncg
Snhg11
Tubb2a
Ina
Apbb2
Mapt
Tubb4a
Uchl1
Gap43
Gng3
Stmn3
Stmn4
Rtn1
Dpysl2
Dpysl3
Cd200
Syt4
Nefl
Pou4f1
Nrp1
Pou6f2
Rgs4
Islr2
Scn9a
Nell2
Hmgcs1
Ebf1
Snca
Fdps
Acat2
Nefm
Mgst3
Atp2b1
Dynll1
Tuba1b
Fgf15
Qk
Btg1
Zeb2
Car14
Mgarp
Cdk4
Hmga1
Mrps33
Mdk
Rps14
Rpl18a
Rps18
Rpl37a
Rorb
Dkk3
Rps3
Sox11
Sox4
Mir124a−1hg
Nupr1
Ndrg1
Pax6
Chd4
Onecut2
Ldhb
Tfap2b
Nrxn3
Mir124−2hg
St18
Gal
Btg2
Ppib
Krt73
Chodl
Igfbp5
Rassf4
Srrm4
N4 node
Sncg
mt−Nd4l
mt−Nd2
mt−Nd1
mt−Co1
mt−Nd3
mt−Atp6
mt−Co2
mt−Nd4
mt−Co3
mt−Cytb
Dynll1
Rps14
Rps18
Rpl37
Rpl6
Rpl26
Rpl23a
Rpl32
Rps6
Rps5
Rps27a
Rps27
Rps8
Rps16
Rpl17
Rps26
Rpl39
Ppia
Rps19
Rpl41
Rps7
Rpl10
Rpl35
Rpl7
Rps11
Rpl37a
Rps28
Rps23
Rpl13a
Rpl36
Tmsb4x
Cfl1
Sepw1
Tmsb10
Stmn1
Ptma
H3f3b
H3f3a
N5 node

### Table S3

| Gene Name | Wave Number |
|---|---|
| Igfbp5 | 1 |
| Hes6 | 1 |
| Nckap5 | 1 |
| Btg2 | 1 |
| Lbr | 1 |
| Gad2 | 1 |
| Dapl1 | 1 |
| Hat1 | 1 |
| Dlx1 | 1 |
| Mdk | 1 |
| Cbfa2t2 | 1 |
| Nkain4 | 1 |
| Pcmtd2 | 1 |
| Itm2a | 1 |
| Hmgn5 | 1 |
| Tpd52 | 1 |
| Car2 | 1 |
| Mgarp | 1 |
| Psip1 | 1 |
| Slc2a1 | 1 |
| Hmgn2 | 1 |
| Miat | 1 |
| Srrm4 | 1 |
| Chchd2 | 1 |
| Cald1 | 1 |
| Gadd45a | 1 |
| Mcm2 | 1 |
| Rassf4 | 1 |
| Tspan9 | 1 |
| Ccer2 | 1 |
| Serpinh1 | 1 |
| Dkk3 | 1 |
| Fgf15 | 1 |
| Ccnd1 | 1 |
| Ddit4 | 1 |
| Btg1 | 1 |
| Mars | 1 |
| Lsm4 | 1 |
| Pou4f2 | 1 |
| Rpl13 | 1 |
| Mir124a-1hg | 1 |
| Elavl3 | 1 |
| Chrna3 | 1 |
| Chrnb4 | 1 |
| Tipin | 1 |
| Ppib | 1 |
| Hmgn3 | 1 |
| Manf | 1 |
| Selm | 1 |
| H2afv | 1 |
| Mfap4 | 1 |
| Nlk | 1 |
| Fam58b | 1 |
| Hist1h1e | 1 |
| Fos | 1 |
| Mfng | 1 |
| Lgals1 | 1 |
| Mcm4 | 1 |
| Ranbp1 | 1 |
| Rps2 | 1 |
| Gm42418 | 1 |
| Lman1 | 1 |
| Dcc | 1 |
| Ctsf | 1 |
| mt-Co2 | 1 |

| Gene Name | Wave Number |
|---|---|
| Tfap2d | 2 |
| Klf7 | 2 |
| Cntn2 | 2 |
| Rnd3 | 2 |
| Neurod1 | 2 |
| Cd59a | 2 |
| Meis2 | 2 |
| Flrt3 | 2 |
| Sulf2 | 2 |
| Dmd | 2 |
| Tsix | 2 |
| Xist | 2 |
| Prdx4 | 2 |
| Zfhx4 | 2 |
| Smc4 | 2 |
| Crabp2 | 2 |
| Slc39a1 | 2 |
| Rbm15 | 2 |
| H2afz | 2 |
| Igfbpl1 | 2 |
| Smc2 | 2 |
| Elavl4 | 2 |
| Slit2 | 2 |
| Rbpj | 2 |
| Hmgb1 | 2 |
| Ptn | 2 |
| Ldhb | 2 |
| Svip | 2 |
| Nr2f2 | 2 |
| Abhd2 | 2 |
| Mki67 | 2 |
| Dpysl4 | 2 |
| Ifitm2 | 2 |
| Cited2 | 2 |
| Arg1 | 2 |
| Akap7 | 2 |
| Foxo3 | 2 |
| Arid5b | 2 |
| Hsp90b1 | 2 |
| Myo16 | 2 |
| Hmgb2 | 2 |
| Tubb3 | 2 |
| Rpgrip1 | 2 |
| Pou4f1 | 2 |
| Spry2 | 2 |
| Ramp3 | 2 |
| Hist3h2ba | 2 |
| Hist3h2a | 2 |
| Alkbh5 | 2 |
| Kdm6b | 2 |
| Sox4 | 2 |
| Dek | 2 |
| Cks2 | 2 |
| Cplx2 | 2 |
| Isl1 | 2 |
| Sox11 | 2 |
| Timmdc1 | 2 |
| Pim1 | 2 |
| AY036118 | 2 |
| Gal | 2 |
| Slc18a2 | 2 |

| Gene Name | Wave Number |
|---|---|
| Mstn | 3 |
| Cntnap5a | 3 |
| Rxrg | 3 |
| Tshz2 | 3 |
| Tmem27 | 3 |
| Phc3 | 3 |
| Ntng1 | 3 |
| Gm30382 | 3 |
| Cnr1 | 3 |
| Rpa2 | 3 |
| Sema3a | 3 |
| Lcorl | 3 |
| Pcdh7 | 3 |
| Apbb2 | 3 |
| A930011G23Rik | 3 |
| Gigyf1 | 3 |
| Tecpr1 | 3 |
| Cttnbp2 | 3 |
| Nav2 | 3 |
| Slc17a6 | 3 |
| Prc1 | 3 |
| D430042O09Rik | 3 |
| Hsf2 | 3 |
| Anks1b | 3 |
| Calb2 | 3 |
| Cdh13 | 3 |
| Dnajc9 | 3 |
| 3632451O06Rik | 3 |
| Pcdh17 | 3 |
| Crabp1 | 3 |
| Tmem108 | 3 |
| Bsn | 3 |
| Snrk | 3 |
| Fstl4 | 3 |
| Car10 | 3 |
| Pde4d | 3 |
| Ckb | 3 |
| Sybu | 3 |
| Nell2 | 3 |
| Kmt2d | 3 |
| Vps8 | 3 |
| Lsamp | 3 |
| Cadm2 | 3 |
| Pla2g7 | 3 |
| Syt4 | 3 |
| Onecut2 | 3 |
| Pdcd4 | 3 |
| mt-Nd1 | 3 |
| mt-Co1 | 3 |
| mt-Nd3 | 3 |

| Gene Name | Wave Number |
|---|---|
| Map2 | 4 |
| Kif1a | 4 |
| St8sia4 | 4 |
| Mgst3 | 4 |
| Tubb4b | 4 |
| Kif5c | 4 |
| Scn3a | 4 |
| Scn9a | 4 |
| Syt13 | 4 |
| Slc24a5 | 4 |
| Myef2 | 4 |
| Pcna | 4 |
| Chgb | 4 |
| Snap25 | 4 |
| Stmn3 | 4 |
| Dcx | 4 |
| Stmn2 | 4 |
| Tnik | 4 |
| Fstl5 | 4 |
| S100a10 | 4 |
| Mllt11 | 4 |
| Igsf3 | 4 |
| Nhlh2 | 4 |
| Elavl2 | 4 |
| Stmn1 | 4 |
| Gnb1 | 4 |
| Agrn | 4 |
| Cyp51 | 4 |
| Insig1 | 4 |
| Tyms | 4 |
| Crmp1 | 4 |
| Nsg1 | 4 |
| Uchl1 | 4 |
| Rufy3 | 4 |
| Asns | 4 |
| Snca | 4 |
| Iqsec1 | 4 |
| Zfp444 | 4 |
| Lgals7 | 4 |
| Sema4b | 4 |
| Arl6ip1 | 4 |
| Sbk1 | 4 |
| Mcmbp | 4 |
| Akap12 | 4 |
| Cd24a | 4 |
| Pcbp3 | 4 |
| Apc2 | 4 |
| Atp2b1 | 4 |
| Dusp26 | 4 |
| Msmo1 | 4 |
| Ncan | 4 |
| Nrp1 | 4 |
| Sncg | 4 |
| Atp8a2 | 4 |
| Stmn4 | 4 |
| Dpysl2 | 4 |
| Nefm | 4 |
| Nefl | 4 |
| Klhl1 | 4 |
| Ncam1 | 4 |
| Islr2 | 4 |
| Scg3 | 4 |
| Elovl4 | 4 |
| Clasp2 | 4 |
| Ebf1 | 4 |
| Mapt | 4 |
| Srsf2 | 4 |
| Actg1 | 4 |
| Pfkp | 4 |
| Pou6f2 | 4 |
| Zscan26 | 4 |
| Tubb2a | 4 |
| Tubb2b | 4 |
| Cltb | 4 |
| Spock1 | 4 |
| Hmgcr | 4 |
| Map1b | 4 |
| Plk2 | 4 |
| Hmgcs1 | 4 |
| Kidins220 | 4 |
| Rtn1 | 4 |
| Lgmn | 4 |
| Chga | 4 |
| Meg3 | 4 |
| Ndrg1 | 4 |
| Cntn1 | 4 |
| Rbfox1 | 4 |
| Rfc4 | 4 |
| Gap43 | 4 |
| Tagln3 | 4 |
| Robo2 | 4 |
| Wtap | 4 |
| Abcg1 | 4 |
| Khsrp | 4 |
| Tubb4a | 4 |
| Lbh | 4 |
| Dpysl3 | 4 |
| Gng3 | 4 |
| Ina | 4 |

| Gene Name | Wave Number |
|---|---|
| Resp18 | 5 |
| Hjurp | 5 |
| Pam | 5 |
| 2310035C23Rik | 5 |
| Dnm3 | 5 |
| P2rx3 | 5 |
| Mpped2 | 5 |
| Sema6d | 5 |
| Pcsk2 | 5 |
| Nnat | 5 |
| Snhg11 | 5 |
| Syp | 5 |
| Il1rapl1 | 5 |
| Pak3 | 5 |
| Serpini1 | 5 |
| Gria2 | 5 |
| Ank2 | 5 |
| Runx1t1 | 5 |
| Jun | 5 |
| Pgm2 | 5 |
| Tmem243 | 5 |
| Magi2 | 5 |
| Sez6l | 5 |
| Cux2 | 5 |
| Kdelr2 | 5 |
| Tac1 | 5 |
| Ccdc136 | 5 |
| Cntnap2 | 5 |
| Mthfd2 | 5 |
| Peg3 | 5 |
| Lig1 | 5 |
| Ube3a | 5 |
| Ndn | 5 |
| Pold3 | 5 |
| Sez6l2 | 5 |
| Cdkn1c | 5 |
| Rmst | 5 |
| Dusp6 | 5 |
| Syt1 | 5 |
| Gm26632 | 5 |
| March1 | 5 |
| Zfhx3 | 5 |
| Tsc22d1 | 5 |
| Pcdh9 | 5 |
| Ntm | 5 |
| Aplp2 | 5 |
| AI593442 | 5 |
| Gm16141 | 5 |
| Sept4 | 5 |
| Dusp3 | 5 |
| Kpna2 | 5 |
| Vcan | 5 |
| Pik3r1 | 5 |
| Myt1l | 5 |
| 1810041L15Rik | 5 |
| Cd200 | 5 |
| Cdkn1a | 5 |
| Efna5 | 5 |
| Nrxn1 | 5 |
| Celf4 | 5 |
| 2900055J20Rik | 5 |
| Fth1 | 5 |
| Fam111a | 5 |
| Prune2 | 5 |
| mt-Nd2 | 5 |
| mt-Atp8 | 5 |
| mt-Nd4l | 5 |
| mt-Nd4 | 5 |
| mt-Cytb | 5 |

| Gene Name | Wave Number |
|---|---|
| Col9a1 | 6 |
| 2900060B14Rik | 6 |
| Rgs4 | 6 |
| Lrp1b | 6 |
| Zeb2 | 6 |
| Rprm | 6 |
| Celf3 | 6 |
| Vax2os | 6 |
| Chl1 | 6 |
| Pou2f2 | 6 |
| D930028M14Rik | 6 |
| Fxyd7 | 6 |
| Synm | 6 |
| Dlg2 | 6 |
| Cacng3 | 6 |
| Igf1 | 6 |
| Kitl | 6 |
| Mgat4c | 6 |
| Prim1 | 6 |
| Zmat4 | 6 |
| Irx6 | 6 |
| Synpr | 6 |
| Pnoc | 6 |
| Onecut1 | 6 |
| Eomes | 6 |
| Acsl6 | 6 |
| Hist1h4d | 6 |
| Glrx | 6 |
| Mast4 | 6 |
| Id2 | 6 |
| Lrfn5 | 6 |
| Six6 | 6 |
| Nrxn3 | 6 |
| Cpne8 | 6 |
| Abat | 6 |
| Glo1 | 6 |
| Gm26917 | 6 |
| Slc3a2 | 6 |
| Tle4 | 6 |

### Table S4

Youngest RGCs (peripheral retina)

| Gene Name | p_val | avg_logFC | p_val_adj |
|---|---|---|---|
| *Mdk* | 1.28E-95 | 2.0447521 | 1.94E-91 |
| *Gal* | 0.905291 | 1.3184257 | 1 |
| *Mfap4* | 1.71E-84 | 1.1971159 | 2.59E-80 |
| *Rassf4* | 1.38E-51 | 1.1322253 | 2.10E-47 |
| *Btg2* | 1.50E-34 | 1.130331 | 2.28E-30 |
| *Mfng* | 7.40E-61 | 1.0969087 | 1.12E-56 |
| *Dlx1* | 1.55E-36 | 1.0702747 | 2.35E-32 |
| *Tead2* | 1.28E-49 | 1.0351948 | 1.94E-45 |
| *Miat* | 3.91E-36 | 0.972397 | 5.93E-32 |
| *Serpinh1* | 1.77E-23 | 0.9567684 | 2.69E-19 |
| *Cdk4* | 5.65E-66 | 0.939143 | 8.58E-62 |
| *Hmgn3* | 3.11E-18 | 0.9187488 | 4.72E-14 |
| *H2afv* | 2.47E-09 | 0.833622 | 3.74E-05 |
| *Prdx1* | 5.98E-34 | 0.817292 | 9.07E-30 |
| *Mfap2* | 5.33E-25 | 0.8124765 | 8.09E-21 |
| *Gadd45a* | 8.35E-36 | 0.7833935 | 1.27E-31 |
| *Hes6* | 4.82E-14 | 0.7814738 | 7.31E-10 |
| *Rps18* | 1.01E-65 | 0.7662706 | 1.53E-61 |
| *Mrps33* | 1.81E-28 | 0.7621987 | 2.74E-24 |
| *Oaz1* | 2.63E-64 | 0.7519175 | 3.99E-60 |
| *Atp6v0e* | 5.13E-27 | 0.7472882 | 7.78E-23 |
| *Srrm4* | 6.65E-23 | 0.7270745 | 1.01E-18 |
| *Gad2* | 3.63E-24 | 0.7265528 | 5.51E-20 |
| *Acp1* | 4.35E-26 | 0.7156413 | 6.60E-22 |
| *Rpl13* | 8.44E-11 | 0.7115311 | 1.28E-06 |
| *Pou4f2* | 2.68E-17 | 0.7068149 | 4.07E-13 |
| *Cbx1* | 3.30E-23 | 0.7067618 | 5.01E-19 |
| *Hmgn2* | 1.05E-20 | 0.6995669 | 1.60E-16 |
| *Grb10* | 4.35E-17 | 0.6859179 | 6.61E-13 |
| *Hnrnpm* | 7.08E-27 | 0.6817614 | 1.07E-22 |

Young RGCS

| Gene Name | p_val | avg_logFC | p_val_adj |
|---|---|---|---|
| *Ccer2* | 1.73E-78 | 1.241453608 | 2.62E-74 |
| *Cntn2* | 4.58E-116 | 1.186168024 | 6.95E-112 |
| *Ppp1r17* | 3.86E-46 | 0.70269425 | 5.86E-42 |
| *Rassf4* | 5.48E-56 | 0.691453756 | 8.32E-52 |
| *Ptma* | 6.09E-130 | 0.690823639 | 9.24E-126 |
| *Igfbp5* | 4.59E-43 | 0.644639818 | 6.97E-39 |
| *Isl1* | 1.77E-72 | 0.629211751 | 2.68E-68 |
| *Skil* | 2.31E-28 | 0.609949218 | 3.51E-24 |
| *E130114P18Rik* | 8.21E-64 | 0.60127407 | 1.25E-59 |
| *Akap7* | 7.75E-23 | 0.596430747 | 1.18E-18 |
| *Elavl4* | 2.78E-63 | 0.572824779 | 4.21E-59 |
| *Gm17750* | 1.78E-42 | 0.529996271 | 2.69E-38 |
| *Chrna3* | 1.17E-25 | 0.526048582 | 1.77E-21 |
| *Adgrg1* | 2.59E-34 | 0.514632025 | 3.94E-30 |
| *Hist3h2a* | 6.92E-27 | 0.513544167 | 1.05E-22 |
| *Peli2* | 4.33E-36 | 0.499921329 | 6.57E-32 |
| *Pou3f1* | 3.68E-41 | 0.491059512 | 5.59E-37 |
| *Sox4* | 1.13E-35 | 0.490708908 | 1.72E-31 |
| *Tub* | 1.61E-32 | 0.486962931 | 2.44E-28 |
| *Nkain4* | 8.99E-25 | 0.484487803 | 1.36E-20 |
| *Tcf4* | 1.81E-30 | 0.471691602 | 2.74E-26 |
| *Dapl1* | 3.68E-24 | 0.46353841 | 5.58E-20 |
| *Thsd7b* | 1.38E-26 | 0.445995086 | 2.10E-22 |
| *Slc16a3* | 5.25E-23 | 0.444327817 | 7.97E-19 |
| *Cplx2* | 3.68E-21 | 0.433504046 | 5.59E-17 |
| *Epb41l3* | 8.09E-21 | 0.428120796 | 1.23E-16 |
| *Pkm* | 4.67E-45 | 0.427276194 | 7.09E-41 |
| *Insc* | 1.14E-29 | 0.410294191 | 1.72E-25 |
| *Cdk2ap1* | 6.97E-22 | 0.410196238 | 1.06E-17 |
| *Igfbpl1* | 4.21E-18 | 0.409705374 | 6.39E-14 |

Old RGCs

| Gene Name | p_val | avg_logFC | p_val_adj |
|---|---|---|---|
| *Apbb2* | 1.56E-73 | 0.9345861 | 2.36E-69 |
| *Mstn* | 2.66E-49 | 0.6422384 | 4.04E-45 |
| *Pou4f1* | 3.57E-34 | 0.6281971 | 5.42E-30 |
| *Crabp1* | 7.75E-06 | 0.6142701 | 0.117549 |
| *Zfhx3* | 4.01E-37 | 0.5370911 | 6.09E-33 |
| *Nefl* | 2.63E-20 | 0.5285437 | 4.00E-16 |
| *Nrn1* | 9.55E-40 | 0.5049236 | 1.45E-35 |
| *Calb2* | 4.53E-21 | 0.5017902 | 6.88E-17 |
| *Ppp2r2b* | 6.16E-32 | 0.4898083 | 9.35E-28 |
| *Nefm* | 4.32E-27 | 0.4567671 | 6.55E-23 |
| *S100a10* | 1.35E-38 | 0.4317273 | 2.04E-34 |
| *Txnrd3* | 1.46E-25 | 0.4280208 | 2.21E-21 |
| *Gap43* | 7.37E-53 | 0.3992509 | 1.12E-48 |
| *Syt4* | 6.55E-29 | 0.3957606 | 9.93E-25 |
| *Stmn4* | 8.85E-29 | 0.3874013 | 1.34E-24 |
| *Islr2* | 2.44E-25 | 0.3868132 | 3.70E-21 |
| *Sncg* | 8.13E-21 | 0.3761556 | 1.23E-16 |
| *Ebf3* | 2.92E-20 | 0.3605836 | 4.44E-16 |
| *Mgst3* | 2.92E-25 | 0.3533699 | 4.43E-21 |
| *Rbp1* | 7.33E-16 | 0.3525898 | 1.11E-11 |
| *Snap25* | 1.08E-23 | 0.3525138 | 1.63E-19 |
| *Ttc28* | 9.91E-20 | 0.3511867 | 1.50E-15 |
| *1700019D03Rik* | 1.13E-24 | 0.3475516 | 1.72E-20 |
| *Uchl1* | 4.31E-27 | 0.3442002 | 6.55E-23 |
| *Nell2* | 6.41E-20 | 0.3394373 | 9.72E-16 |
| *Gm30382* | 9.25E-17 | 0.3341243 | 1.40E-12 |
| *Acat2* | 1.89E-18 | 0.3320028 | 2.86E-14 |
| *Khdrbs2* | 2.43E-16 | 0.3292614 | 3.69E-12 |
| *Ppp1r1c* | 1.62E-20 | 0.3172344 | 2.46E-16 |
| *Gpr37* | 3.77E-21 | 0.3114471 | 5.72E-17 |

Oldest RGCs (central retina)

| Gene Name | p_val | avg_logFC | p_val_adj |
|---|---|---|---|
| *Snhg11* | 1.79E-114 | 1.365474193 | 2.71E-110 |
| *Igf1* | 1.68E-65 | 1.34349882 | 2.55E-61 |
| *Meg3* | 3.82E-104 | 1.089751573 | 5.79E-100 |
| *Mast4* | 2.32E-68 | 1.086630225 | 3.52E-64 |
| *Celf4* | 5.97E-108 | 0.95833074 | 9.06E-104 |
| *Synm* | 1.38E-83 | 0.953486758 | 2.10E-79 |
| *Pcdh9* | 3.51E-100 | 0.953445391 | 5.33E-96 |
| *Fxyd7* | 9.33E-95 | 0.942849091 | 1.42E-90 |
| *Synpr* | 5.55E-70 | 0.937301446 | 8.43E-66 |
| *Kitl* | 8.81E-73 | 0.910637823 | 1.34E-68 |
| *Ntm* | 2.94E-86 | 0.906603729 | 4.46E-82 |
| *Myt1l* | 2.88E-101 | 0.901995121 | 4.37E-97 |
| *Tubb2a* | 1.27E-110 | 0.863992223 | 1.93E-106 |
| *Eomes* | 1.95E-60 | 0.819124315 | 2.97E-56 |
| *Lrfn5* | 6.62E-80 | 0.738675402 | 1.01E-75 |
| *Rbms3* | 7.22E-74 | 0.719680318 | 1.10E-69 |
| *Cntnap2* | 1.13E-66 | 0.715428154 | 1.71E-62 |
| *Syt4* | 6.64E-57 | 0.703630893 | 1.01E-52 |
| *Rprm* | 1.39E-36 | 0.697665498 | 2.12E-32 |
| *D930028M14Rik* | 6.09E-74 | 0.681902252 | 9.24E-70 |
| *Mapt* | 1.01E-80 | 0.679990502 | 1.54E-76 |
| *Chl1* | 5.29E-68 | 0.661633097 | 8.03E-64 |
| *Cnr1* | 2.61E-56 | 0.646538864 | 3.96E-52 |
| *Ncam1* | 1.03E-49 | 0.638042873 | 1.57E-45 |
| *2900011O08Rik* | 6.90E-57 | 0.636688071 | 1.05E-52 |
| *Resp18* | 4.31E-58 | 0.636140301 | 6.54E-54 |
| *Runx1t1* | 4.55E-41 | 0.635924144 | 6.91E-37 |
| *Ube3a* | 3.28E-47 | 0.623037196 | 4.98E-43 |
| *Pou6f2* | 1.09E-48 | 0.607352764 | 1.66E-44 |
| *Car10* | 3.59E-49 | 0.606612349 | 5.45E-45 |

*Table S5*

Orientation markers

| Dorsal | Ventral | Temporal | Nasal |
|---|---|---|---|
| *Efnb1* | *Epha6* | *Epha5* | *Efna5* |
| *Efnb2* | *Ephb2* | *Epha6* | *FoxG1* |
| *Sox12* | *Ephb3* | *Ephb1* | |
| *Tbx5* | *Ephb4* | *EphA3* | |
| *Fstl4* | *Vax2* | *Foxd1* | |
